# Acetogenic methane-carbon monoxide comproportionation: an exergonic but unobserved microbial metabolism

**DOI:** 10.64898/2026.01.08.698499

**Authors:** Heidi S. Aronson, William D. Leavitt, Douglas E. LaRowe

## Abstract

Microbial metabolism relies on redox reactions that exploit chemical disequilibria. While aerobic carbon oxidation, carbon fixation, and fermentation are well studied, the broader space of anaerobic carbon redox reactions remains underexplored. In this study, carbon comproportionation, or reverse fermentation, reactions are identified as a previously unrecognized and potentially favorable class of microbial carbon redox transformations. Particular attention is given to the reaction between methane (CH4) and carbon monoxide (CO) to form acetate, a reaction that has not previously been evaluated despite the widespread occurrence of CH4 and CO in anoxic systems. Gibbs energies (Δ*G_r_*) for this reaction were calculated across broad ranges of temperature, pH, and dissolved CH4 and CO concentrations using measured physicochemical data from a wide variety of environmental systems. We show that *acetogenic* CH4-CO *comproportionation* is exergonic in all environments where both substrates were detected. The most favorable energetic conditions occur at high pH, low temperature, and high reactant concentrations, consistent with cool serpentinizing systems. In several settings, the calculated Gibbs energy yields and energy densities overlap or exceed known anaerobic metabolisms involving CH4, CO, and acetate. These results demonstrate that acetogenic CH4-CO comproportionation can support microbial energy conservation in a variety of settings. To determine if this metabolism could have operated on early Earth or Mars, modeled fluid compositions show that this reaction is also exergonic under plausible physicochemical regimes. This work broadens the suite of possible microbial energy metabolisms and provides testable criteria for evaluating carbon-based catabolic reactions on Earth and on other planetary bodies.

**Plain Language Summary:** Microorganisms obtain energy by catalyzing chemical reactions in their environment. The energy available from a reaction can be quantified using Gibbs energies of reaction (Δ*G_r_*). When Δ*G_r_* < 0, energy is released that microorganisms can use to build biomass and carry out other activities. In this study, we predicted a new energy-yielding reaction that could potentially support microbial life. In this reaction, methane (CH4) is oxidized using carbon monoxide (CO) to produce acetate. Using thermodynamic calculations and measured geochemical data from natural environments, we show that this reaction can release usable energy under a wide range of conditions, including continental serpentinizing systems, the deep continental and marine subsurface, and geothermal springs. We also predict that this reaction could support life under plausible early Earth conditions and in modeled Martian fluids. Together, these observations identify the reaction of CH4 and CO to form acetate as a potentially viable microbial energy source in anoxic environments on Earth and other planetary bodies.

## Introduction

Some microorganisms obtain energy and carbon from chemoautotrophic and chemoorganotrophic metabolic reactions. Over the past half century, first principles thermodynamic calculations (theory) have been utilized to predict microbial catabolic reactions prior to their discovery in nature (empirical evidence). If a redox reaction is sufficiently exergonic (energy-yielding), it can, in principle, support microbial life. This thermodynamics-guided framework has identified catabolic reactions that are now recognized as major contributors to global biogeochemical cycling. Notable examples include anaerobic ammonia oxidation with nitrite or nitrate (anammox) (Broda, 1977; Van de Graaf et al., 1995), which accounts for an estimated 30-50% of oceanic nitrogen loss (Hutchins & Capone, 2022), and the anaerobic oxidation of methane with sulfate (AOM) (Barnes & Goldberg, 1976; Boetius et al., 2000; Hinrichs et al., 1999; Orphan et al., 2001), which consumes 85-300 Tg of methane annually (Knittel & Boetius, 2009). The prediction of complete ammonia oxidation (comammox) likewise preceded its discovery in nature (Costa et al., 2006; Daims et al., 2015; van Kessel et al., 2015). More recently, thermodynamic predictions have suggested additional “missing” metabolisms, including ammonia-driven methanogenesis, a number of redox reactions among Mn-bearing species, and sulfur comproportionation (Amend et al., 2019, 2020; D. E. LaRowe et al., 2021). Indeed, recent experimental work by Aronson and colleagues supports sulfur comproportionation in a strain of *Acidithiobacillus thiooxidans* (Aronson et al., 2025). These studies demonstrate that thermodynamic space still contains plausible yet unrecognized microbial metabolic reactions.

Comproportionation reactions represent one such unexplored class of redox transformations, where two chemical species containing the same element in different oxidation states react to form a product with an intermediate oxidation state. Several known microbial metabolisms follow this pattern, including anammox, which combines nitrite (oxidation state +3) and ammonium (-3) to form N2 (+0) (Van de Graaf et al., 1995), and sulfur comproportionation, which produces elemental sulfur (+0) from sulfate (+6) and sulfide (-2), and for which a putative microbial candidate has been identified. Comproportionation reactions are in essence reverse disproportionation, or fermentation, reactions, in which a compound of intermediate oxidation state acts simultaneously as an electron donor and acceptor to form two new compounds with divergent oxidation state (e.g. glucose fermentation to ethanol and carbon dioxide: C_6_H_12_O_6_ → 2CO_2(aq)_ + 2C_2_H_5_OH, where the comproportionation reaction would be the reverse: 2CO_2(aq)_ + 2C_2_H_5_OH → C_6_H_12_O_6_). Although comproportionation is well established for nitrogen and recently for sulfur, analogous reactions involving carbon have received comparatively little attention, despite carbon’s central role in microbial metabolism (e.g., assimilation, respiration, fermentation and global biogeochemical cycling).

Carbon exists in oxidation state from -4 (e.g. methane) to +4 (e.g. CO2), with many intermediates occupying nominal oxidation states of carbon (NOSC) that can serve as products or substrates in redox transformations (Fig. 1). Numerous combinations of carbon compounds can theoretically undergo carbon comproportionation reactions, and some of these reactions may be sufficiently exergonic to support microbial energy metabolism. However, the breadth of this reaction space complicates the identification of environmentally relevant and physiologically plausible pathways. To evaluate potential carbon comproportionation reactions, we first surveyed a range of candidate reactions spanning a range of carbon oxidation states using calculations of the standard state Gibbs energy (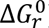) (Fig. 2). This screening step was designed to identify reactions that warrant further consideration under environmentally realistic conditions. From this broader reaction space, we focus on a single carbon comproportionation reaction of particular environmental relevance: the reaction of methane (CH4; NOSC = -4) and carbon monoxide (CO; NOSC = +2) to form acetate (NOSC = 0) (Rxn. 1). This reaction links two C1 carbon compounds that co-occur and are widely cycled in many anoxic environments and produces acetate, a central metabolite in anaerobic carbon metabolism, including acetogenic carbon fixation and fermentation. More broadly, reactions involving carbon monoxide are generally exergonic, yet remain comparatively underexplored relative to other carbon catabolic reactions. Importantly, paired measurements of CO and CH4 are available in a variety of natural sites, enabling quantitative evaluation of this reaction across diverse geochemical settings. This data availability allows acetogenic CH4-CO comproportionation to be assessed not only as a theoretical possibility but also within the geochemical contexts where it may realistically occur. As such, this reaction represents a plausible carbon comproportionation catabolism that could have operated under early Earth conditions, may be viable in extraterrestrial environments, and could be occurring today in certain extreme environments.

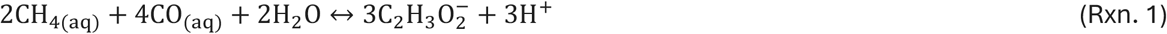

**Figure 1.**
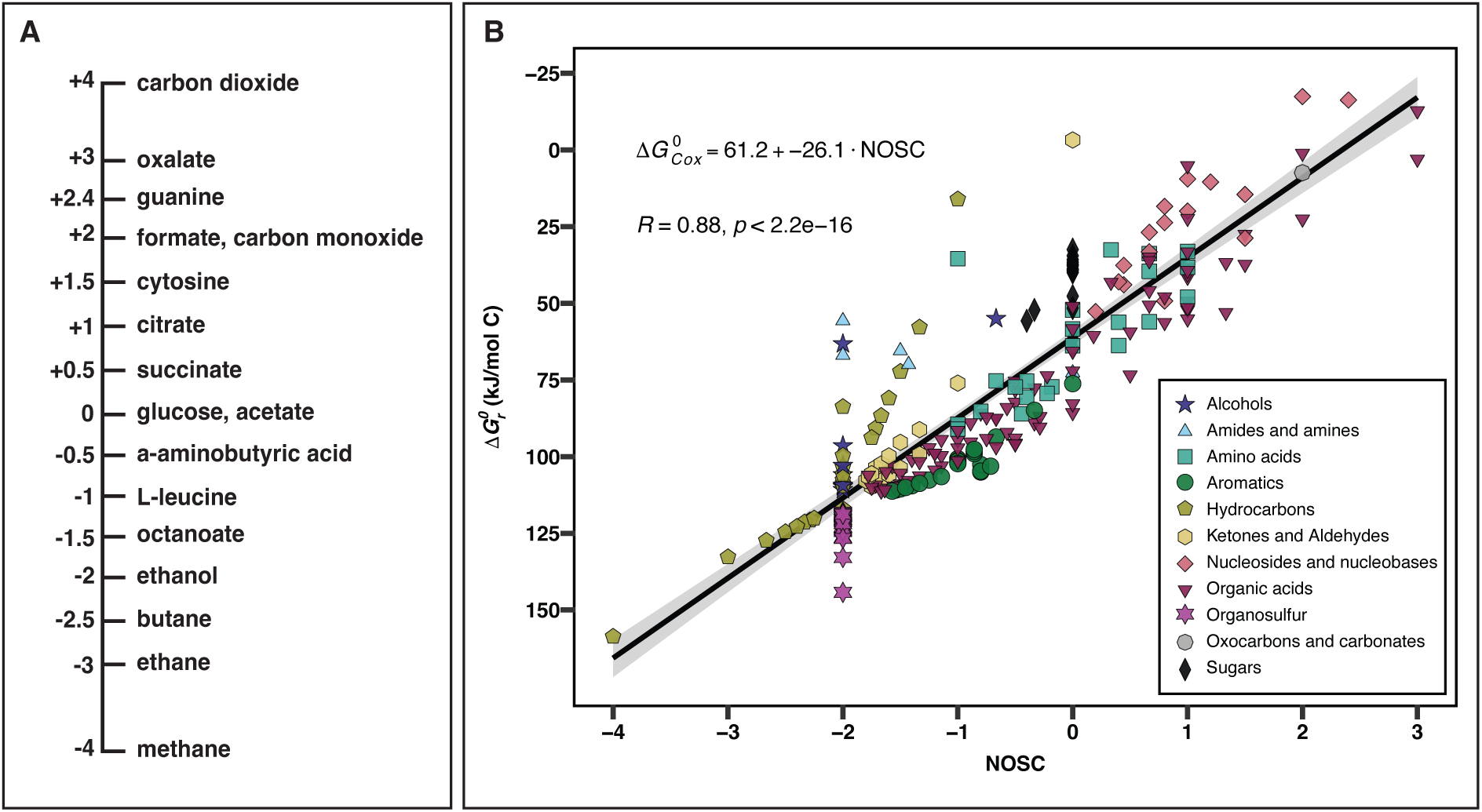
A) Standard state Gibbs energies (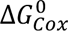) calculated for half reactions describing the complete oxidation of each compound to bicarbonate at 25 °C and 1 bar, as a function of the nominal oxidation states of carbon, or NOSC. The shapes and colors of the symbols correspond to the indicated classes or organic compounds. B) The NOSC of selected carbon compounds.

**Figure 2.**
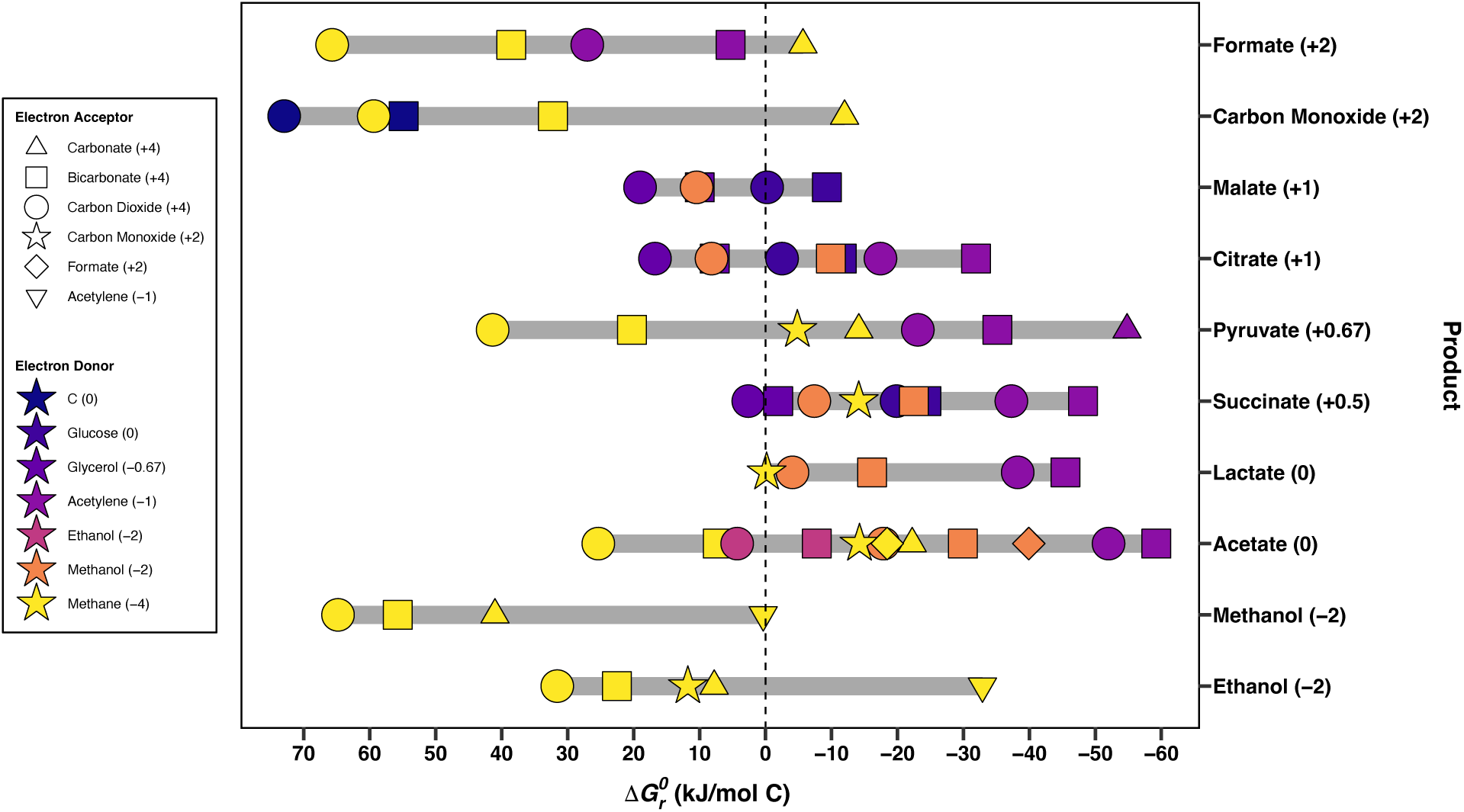
Standard state Gibbs energies, 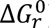 (kJ/mol carbon), at 25 °C for 65 different carbon comproportionation reactions. Each point represents a unique donor-acceptor-product combination. Point color denotes the electron donor, point shape denotes the electron acceptor, and the vertical position corresponds to the product compound (listed along the right side of the plot). For example, the left-most symbol in the top line corresponds to the reaction of 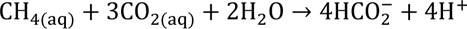. The numbers in parentheses next to each product species refer to the nominal oxidation states of carbon (NOSC) in that compound.

Here, we evaluate the thermodynamic feasibility of acetogenic CH4-CO comproportionation as a potential mode of microbial energy conservation following the approach of Amend et al. (2020) and Aronson et al. (2025) for S-comproportionating metabolisms. We calculate Gibbs energies (Δ*G_r_*) across wide ranges of temperature, pH, and ionic composition. These calculations incorporate measured physicochemical data from several anoxic environment types, including serpentinizing systems, deep continental and marine subsurface fluids and sediments, and geothermal springs, as well as modeled early Earth and Martian conditions. By performing thermodynamic calculations under a variety of actual geochemical environments, this study delineates the natural conditions under which carbon comproportionation reactions may be energetically viable and establishes a framework for assessing their potential role in anoxic ecosystems on Earth, in Earth’s past, and on other planetary bodies.

## Methods

The nominal oxidation state of carbon (NOSC) was calculated for 253 organic compounds from the CHNOSZ database (Dick, 2019) using,

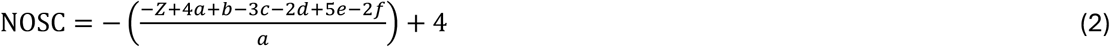

where *Z* is the net charge of the compound, and *a, b, c, d, e,* and *f* correspond to the stoichiometric numbers of the elements C, H, N, O, P, and S in each compound (D. E. LaRowe & Van Cappellen, 2011). A half reaction describing complete oxidation to bicarbonate was written for each compound (Table S1), and the standard state Gibbs energies of carbon oxidation, 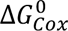, were calculated for each at 25 °C and 1 bar with the revised Helgeson-Kirkham-Flowers (HKF) equations of state (Helgeson et al., 1981, p. 1; Shock & Helgeson, 1988; Tanger & Helgeson, 1988) with the “subcrt” command in the R software package CHNOSZ v1.4.1 (Dick, 2019). Thermodynamic data implemented in CHNOSZ are sourced from the OrganoBioGeoTherm database; original data sources and references are provided in the Supplementary Methods.

The standard state Gibbs energy of reaction, 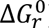, was calculated for 65 carbon comproportionation reactions as described above (Table S2). Electron donors (and their oxidation states) included methane (-4), ethanol (-2), methanol (-2), acetylene (-1), glycerol (-0.67), glucose (0), and elemental carbon as graphite (0). These substrates include widely utilized biological carbon sources (e.g., methane, glucose, methanol, ethanol) and compounds involved in previously proposed or demonstrated comproportionation reactions (Kremp et al., 2018; Lahijani et al., 2015; Lever, 2012; Litty et al., 2022; Steiger et al., 2017). Electron acceptors included CO (+2), formate (+2), and dissolved inorganic carbon (DIC) species CO2/HCO3-/CO32- (+4), in addition to acetylene. Ten product compounds were chosen from physiologically relevant metabolic intermediates: formate (+2), carbon monoxide (+2), malate (+1), citrate (+1), pyruvate (+0.67), succinate (+0.5), lactate (0), acetate (0), methanol (-2), and ethanol (-2). These compounds are common in microbial fermentation reactions, carbon fixation pathways (e.g., Wood-Ljunghdal, reverse TCA), and organic acid metabolism (Ragsdale & Pierce, 2008).

Gibbs energies of reaction, Δ*G_r_*, were calculated for Rxn. 1 with Eqn. 3:

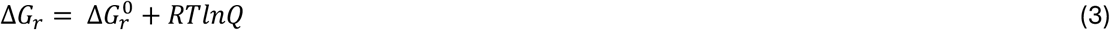

where 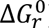 is the standard state Gibbs energy, *R* denotes the universal gas constant, *T* represents to the temperature in Kelvin, and *Q_r_* refers to the reaction quotient. Values of *Q_r_* were calculated using Eqn 4:

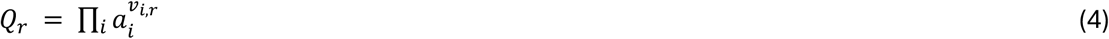

where *a_i_* corresponds to the activity of the *i*th species raised to its stoichiometric coefficient, *a_i,r_*, in the *r*th reaction, which is positive for products and negative for reactants. Activities were calculated with Eqn 5.

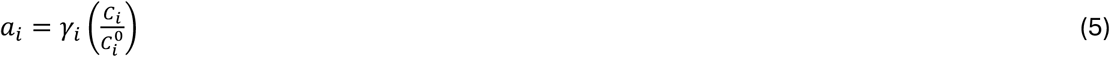

where γ_*i*_ and *C*_*i*_ represent the activity coefficient and the measured concentration of the *i^th^* species, respectively. 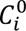 is the concentration of the *i^th^* species under standard state conditions, which is equal to one molal referenced to infinite dilution. Measured concentrations were typically reported in molarity; for activity calculations, these values were treated as equivalent to molality.

Δ*G_i_* was calculated for Rxn. 1 at 1 bar across a range of pH (0-14) and temperature (0-100 °C) with the activities of products and reactants (except for H^+^) set at environmentally relevant activities. The activities of methane (CH4), carbon monoxide (CO), and acetate were set at 10^-5^ (≈10 µM), 10^-9^ (≈1 nM), and 10^6^ (≈1 µM), respectively.

Concentrations of CH4 and CO from a variety of environments were taken from a number of published sources (Boulart et al., 2013; Canovas et al., 2017; Cardace et al., 2015; Cook, Blank, Rietze, et al., 2021; Cook, Blank, Suzuki, et al., 2021; Crespo-Medina et al., 2014; Eickenbusch et al., 2019; Gihring et al., 2006; Howells et al., 2025; Kieft et al., 2005; Lau et al., 2016; Lu et al., 2020; Magnabosco et al., 2016; Mara et al., 2023; Momper et al., 2023; Morrill et al., 2014; Moser et al., 2005; Nyyssönen et al., 2012; Onstott et al., 2009; Osburn et al., 2014, 2019; Qi et al., 2025; Robare, 2025; Sabuda et al., 2021; Twing et al., 2017) (Table S3). For samples in which CO and CH4 were reported as below detection limits, concentrations were assigned as one order of magnitude lower than the analytical detection limit for that compound specified in each study. Within the serpentinite mud samples (Eickenbusch et al., 2019), CH4 was below detection limits (< 0.01 µM) in 10 out of 67 samples and CO was below detection limits (< 0.2 µM) in 16 out of 67 samples. These samples were assigned concentrations of 10^-8^ M CH4 and 2×10^-8^ M CO. Within serpentinizing spring samples, CO was below detection limits (< 10 µM) in 10 out of 88 samples. These samples were assigned concentrations of 10^-7^ M CO. This approach avoids overestimating reactant activities, which would otherwise bias the Δ*G_r_* estimates toward more negative values. The Δ*G_r_* values calculated from these substituted activities should therefore be interpreted as conservative estimates of the potential energy yield of Rxn. 1 under low-CO and low-CH4 conditions. For samples in which acetate was not reported, the acetate concentration was set to 1 µM. This value falls within the range of acetate concentrations measured across many anoxic environments and provides a reasonable estimate for Δ*G_r_* calculations. Activities were calculated from the *in* situ or estimated chemical compositions of each sample using the “ae.speciate” command from the aqueous speciation package AqEquil v0.18.0 (Boyer et al., 2024) (Table S4).

Each 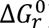 was calculated as described above for 439 natural samples at the reported *in situ* temperatures and pressures. Water pressure at the seafloor was calculated for marine subsurface sediment samples (Supplementary Methods). Geochemical data for hypothetical Mars fluids were estimated using data from (Marlow et al., 2014; Rucker et al., 2023), with atmospheric CO concentrations set at either 1 or 50% (Koyama et al., 2024), for a total of 22 samples. Henry’s law was used to calculate the concentration of aqueous CO from gaseous CO at 7 °C, 9 °C, 14 °C, 15 °C, 24 °C, 48 °C, 65 °C, and 83 °C. Henry’s law constants (mol L^-1^ atm^-1^) for CO were calculated at each temperature using the equilibrium constants of the gas dissolution reaction for each compound at 2.2 bar. Geochemical data for a hypothetical early Earth ocean were estimated by determining the equilibrium activities of aqueous CH4 corresponding to partial pressures ranging from 5×10^-3^ to 1.8×10^-2^ bar and aqueous CO corresponding to partial pressures ranging from 10^-5^ to 10^-2^ bar at temperatures from 0 to 40 °C, pH 7 (Halevy & Bachan, 2017), using the ionic strength of the modern ocean without sulfate, for a total of 164 conditions.

Gibbs energies, Δ*G_r_*, were calculated for Rxn. 1 as described above for 625 samples (439 natural samples and 186 modeled conditions) as described above. Negative values of Δ*G_r_* indicate exergonic (energy-yielding) reactions whereas positive values indicate endergonic (energy-consuming) reactions. To compare Rxn. 1 with known anaerobic microbial metabolic reactions that consume CO or CH4 or produce acetate, Δ*G_r_* was calculated for six additional reactions for each sample under the geochemical parameters described above. These reactions include the water-gas shift reaction, CO oxidation coupled to sulfate reduction, methanogenic CO disproportionation, acetogenic CO disproportionation, methane oxidation coupled to sulfate reduction (AOM), and homoacetogenesis. For sites where sulfate, sulfide, or hydrogen activities were absent, activities of 10^-6^ were assumed.

The reactions evaluated are:

Water gas shift

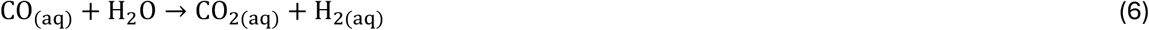

CO oxidation coupled to sulfate reduction

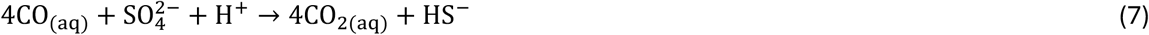

Methanogenic CO disproportionation

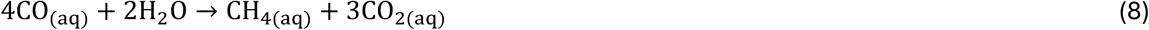

Acetogenic CO disproportionation:

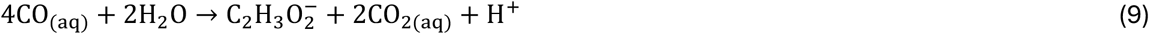

Anaerobic oxidation of methane (AOM)

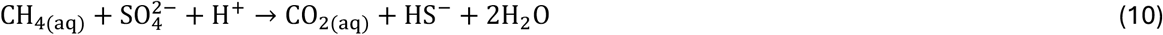

Homoacetogenesis

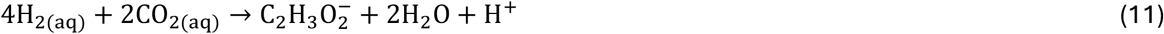

To account for differences in substrate availability among reactions and environments, energy densities (*E_r_*; kJ/kg H2O) were calculated to weight the Gibbs energy by the availability of the limiting reactant:

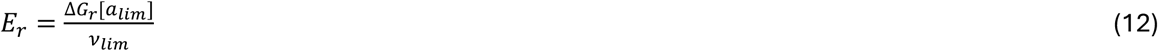

where *a_lim_* is the activity of the limiting reactant and *v*_*lim*_ is its stoichiometric coefficient. Energy densities represent the total accessible energy available per kilogram of water under the specified geochemical conditions. For quantitative comparisons among reactions, Rxn. 1 was treated as a reference reaction, and pairwise differences in the Gibbs energy and energy density were calculated for each sample. Differences in energy density were evaluated using the log-transformed metric Δ*logE*, defined for samples in which both reactions were exergonic as:

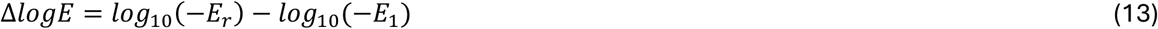

where *E_r_* and *E*_1_are the energy densities of the comparison reaction and Rxn. 1, respectively. Δ*logE* was calculated only for samples in which both reactions are exergonic, as energy densities for endergonic reactions are positive and therefore yield non-real values under the *log*_10_(−*E_r_*) transformation. Negative values of Δ*logE* indicate higher energy density for Rxn. 1, whereas positive values indicate higher energy density for the comparison reaction. For each reaction, the null hypothesis that the median Δ*logE* equals zero was evaluated using a Holm-corrected Wilcoxon signed-rank test. Statistical analyses were conducted across the 439 natural samples only. Statistical analyses were performed using R version 4.5.1. All data and R scripts are available at https://doi.org/10.5281/zenodo.18158636.

## Results

### Thermodynamic potential for carbon comproportionation reactions

Standard-state Gibbs energies of carbon oxidation (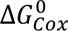) for 253 carbon-containing compounds are shown in Fig. 1A. Examples of NOSC values for common compounds are shown for reference in (Fig. 1B). These compounds span the full range of NOSC values and include alcohols, amides and amines, amino acids, aromatics, hydrocarbons, ketones and aldehydes, nucleosides and nucleobases, organic acids, organosulfur compounds, sugars, oxocarbons, and carbonates. Consistent with earlier work (D. E. LaRowe & Van Cappellen, 2011), 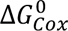 shows an inverse correlation with NOSC (R^2^ = 0.85, p < 2.2×10^-16^). Compounds with higher NOSC values exhibited lower half reaction energies (Fig. 1B).

We present the standard state Gibbs energies 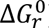, at 25 °C) for 65 carbon comproportionation reactions relevant to microbial carbon transformations (Fig. 2). These reactions span a broad range of NOSC values. Of these the oxidation of acetylene with bicarbonate to produce acetate had the most negative value of 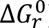 (-59 kJ/mol C), whereas the oxidation of elemental carbon with CO2 to form CO yielded the most positive value (+73 kJ/mol C). Reactions forming pyruvate exhibited the widest range in 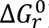. For example, the reaction of methane and CO2 has a standard state Gibbs energy of +41 kJ/mol C while the acetylene-CO32-pair yielded -55 kJ/mol C. By contrast, reactions forming malate showed the narrowest range of 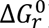, from +19 kJ (mol C) ^-1^ for glycerol-CO2 to +0.4 kJ/mol C for methane-acetylene. Many acetylene-based reactions yielded negative values of 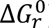 (-17 to -59 kJ/mol C).

The standard-state Gibbs energies of Rxn. 1 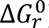) at 1, 1000, and 3000 bar are shown as a function of temperature in Fig. 3. Values of 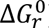 were negative from 0 to 122 °C and became more negative with increasing temperatures, ranging from –15 kJ/mol C at 0 °C and –3 kJ/mol at 122 °C. At each temperature, values of 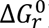 became more negative with increasing pressure. At 0 °C, 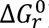 decreased from -15 kJ/mol C at 1 bar to -20 kJ/mol C at 3000 bar, and at 122 °C 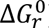 decreased from -3 kJ mol^-1^ at 1 bar to -11 kJ mol^-1^ at 3000 bar. When overall Gibbs energies (Δ*G_r_*) were evaluated at environmentally realistic activities of CH₄ (10 µM), CO (1 nM), and acetate (1 µM), the reaction remained exergonic across broad ranges of pH and temperature (Fig. 4). Under these conditions, values of Δ*G_r_* approached –60 kJ/mol C at high pH and low temperature. The strong pH dependence reflects the consumption of three protons in Rxn. 1; temperature has a secondary influence on energy yield.

**Figure 3.**
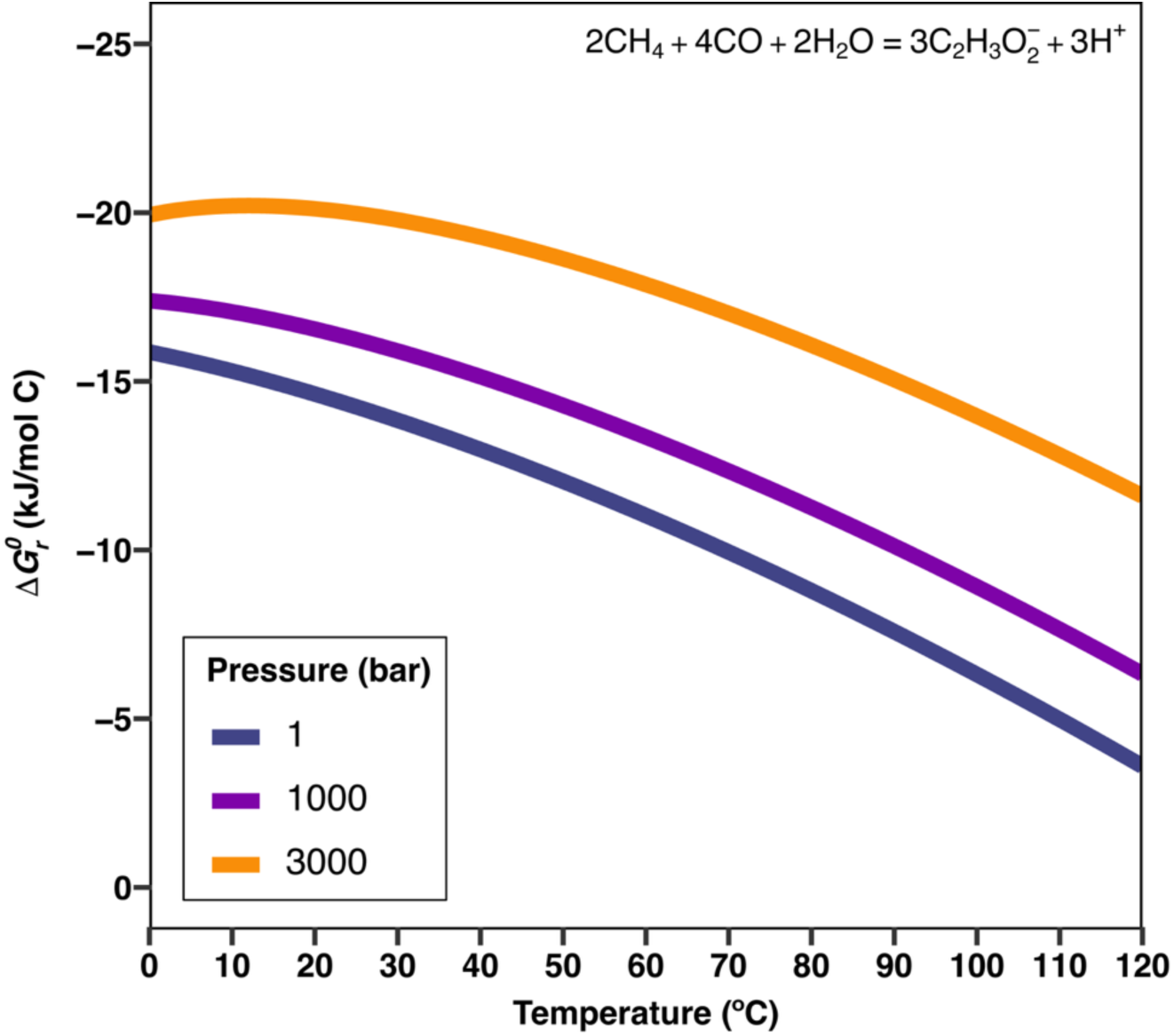
Standard state Gibbs energy, 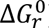 (kJ/mol carbon), for Rxn. 1 as a function of temperature. For temperatures ≥ 100 °C, standard state Gibbs energies were calculated at the saturation vapor pressure of water for each temperature.

**Figure 4.**
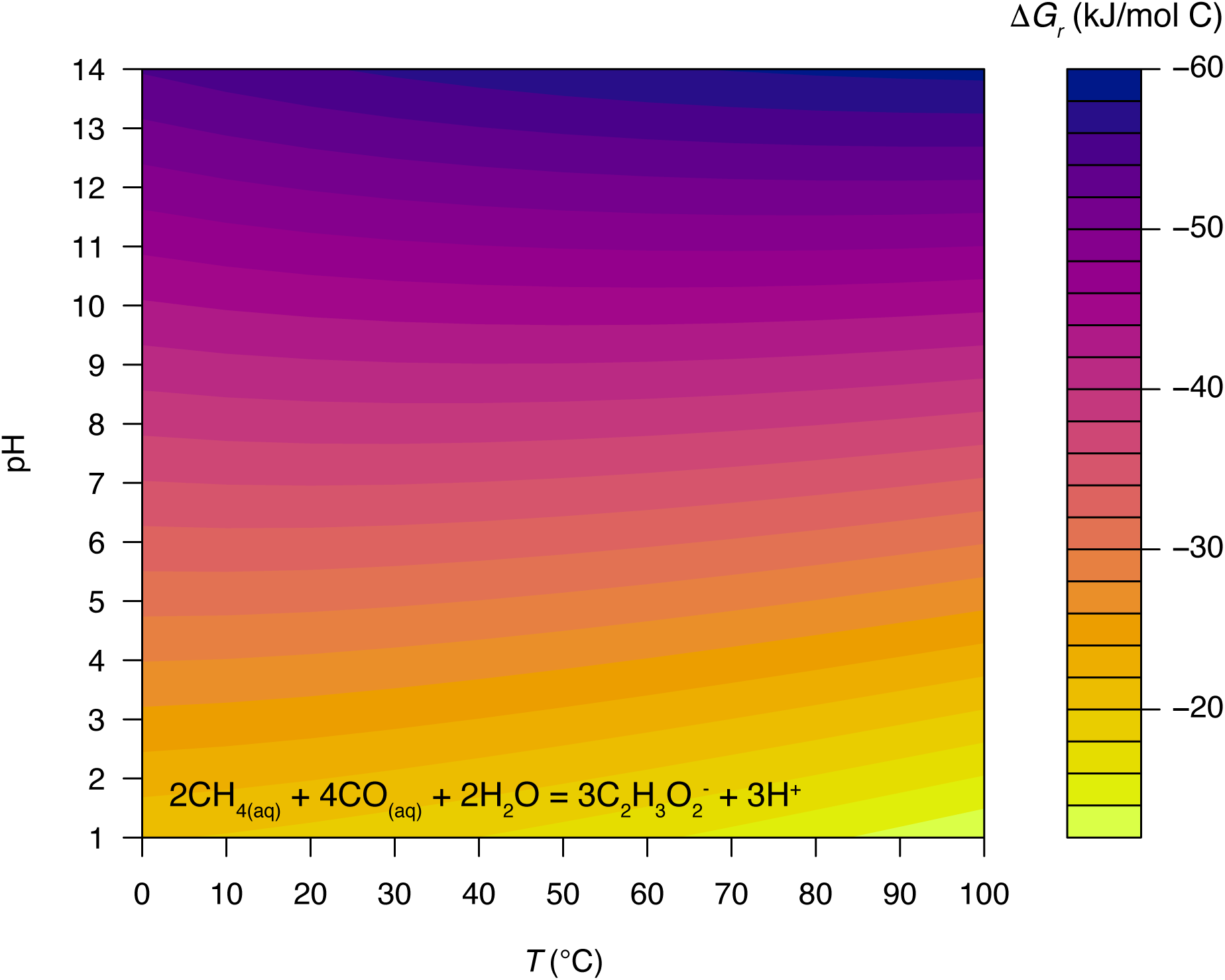
Values of the overall Gibbs energy of reaction, Δ*G_r_* (kJ/mol carbon), for Rxn. 1 as a function of temperature and pH. Activities of acetate and aqueous CH4 and CO were set at10^-6^, 10^-5^, and 10^-9^, respectively.

While not an exhaustive list of datasets, we found that CO and CH4 co-occurred in 25 geochemical datasets that represented 439 natural samples, together with 28 modeled fluids representing early Earth oceans and Martian environments, for a total of 467 scenarios. The energetics of Rxn. 1 across all 467 geochemical scenarios evaluated are plotted in Fig. 5. Across this combined dataset, Δ*G_r_* values normalized by moles of carbon spanned from -42 kJ/mol C to +21 kJ/mol C, with 372 samples revealing negative Gibbs energies (80% of sites). Among the natural samples, pH ranged from 1.8 in springs from Yellowstone National Park to 13.1 at serpentinizing springs at the Tablelands. Temperature ranged from 2 °C in serpentinite muds from the Mariana Forearc to 96 °C in the deep subsurface sediment samples from Guaymas Basin. The deep continental subsurface samples displayed the widest range of CO measurements, with the lowest and highest reported CO concentrations in the entire dataset (0.06 nM at DeMMO to 680 µM at Kloof Mine). The lowest reported CH4 measurement was from DeMMO (0.08 nM) and the highest was from Guaymas Basin (130 mM). To identify the geochemical variables that exert the strongest influence on Δ*G_r_*, correlations were evaluated using only the natural samples (n = 439). Across these samples, pH exhibited a strong linear association with Δ*G_r_* (R^2^ = 0.86, p = 1.96×10^-179^), consistent with the stoichiometric consumption of three protons in Rxn. 1 (Fig. 6A). Temperature also exerted a significant, but secondary association (Fig. 6B) (R^2^ = 0.558, p = 1.63×10^-79^), reflecting the temperature dependence of 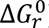 (Fig. 3).

**Figure 5.**
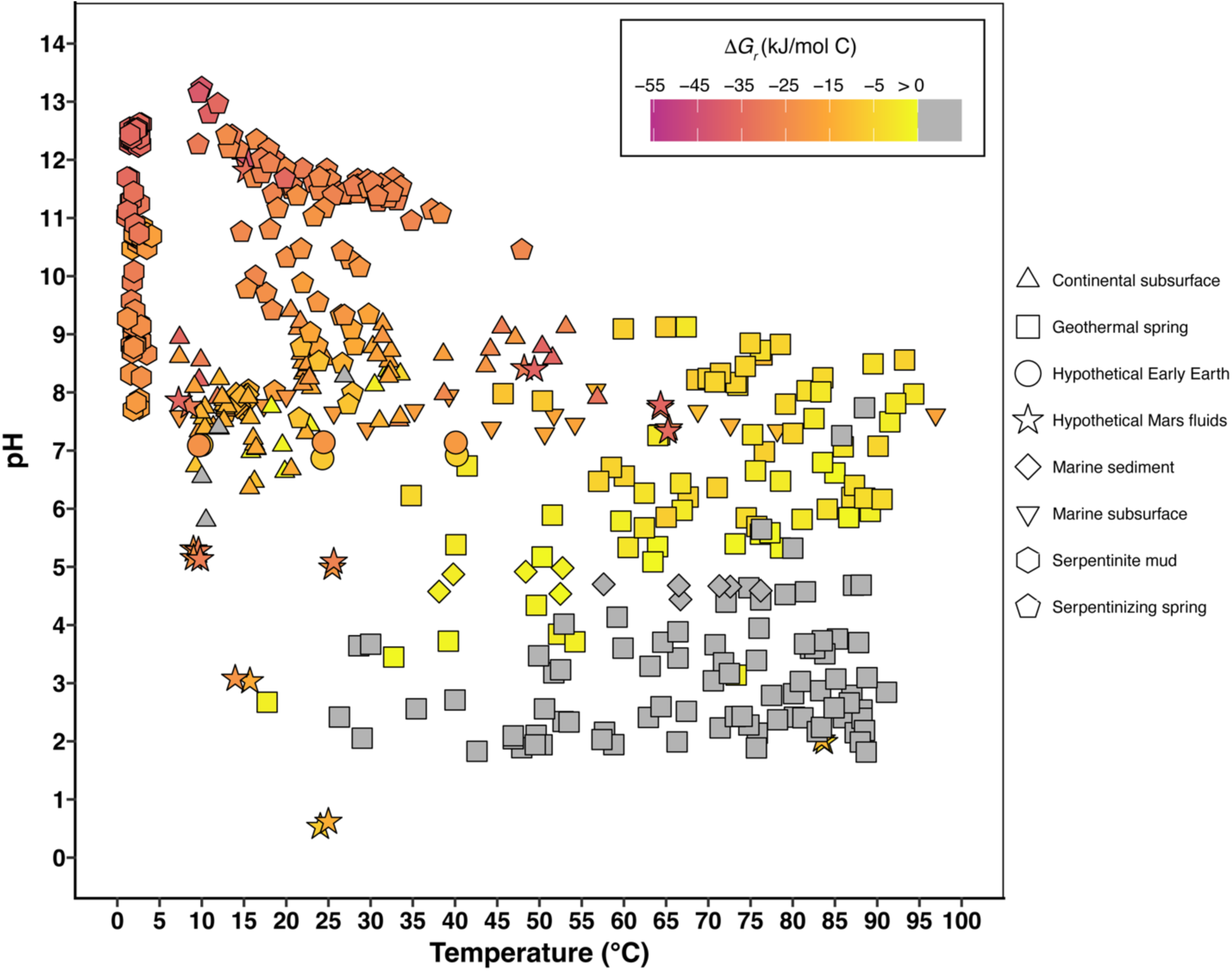
Values of the overall Gibbs energy of reaction, Δ*G_r_* (kJ/mol carbon) for Rxn. 1. Values of Δ*G_r_* are shown as a function of temperature and pH for 467 combinations of temperature, pH, pressure, and activities of CH4, CO, and acetate representing samples from seven anoxic environment types, along with modeled early Earth ocean and Martian fluids. Temperature, pH, pressure and activities of CH4, CO, and acetate were taken directly from *in situ* geochemical data from seven environment types, represented as the shape of the points. Point fill indicates the magnitude of Δ*G_r_* (kJ/mol C); gray points denote samples where Δ*G_r_* > 0.

**Figure 6.**
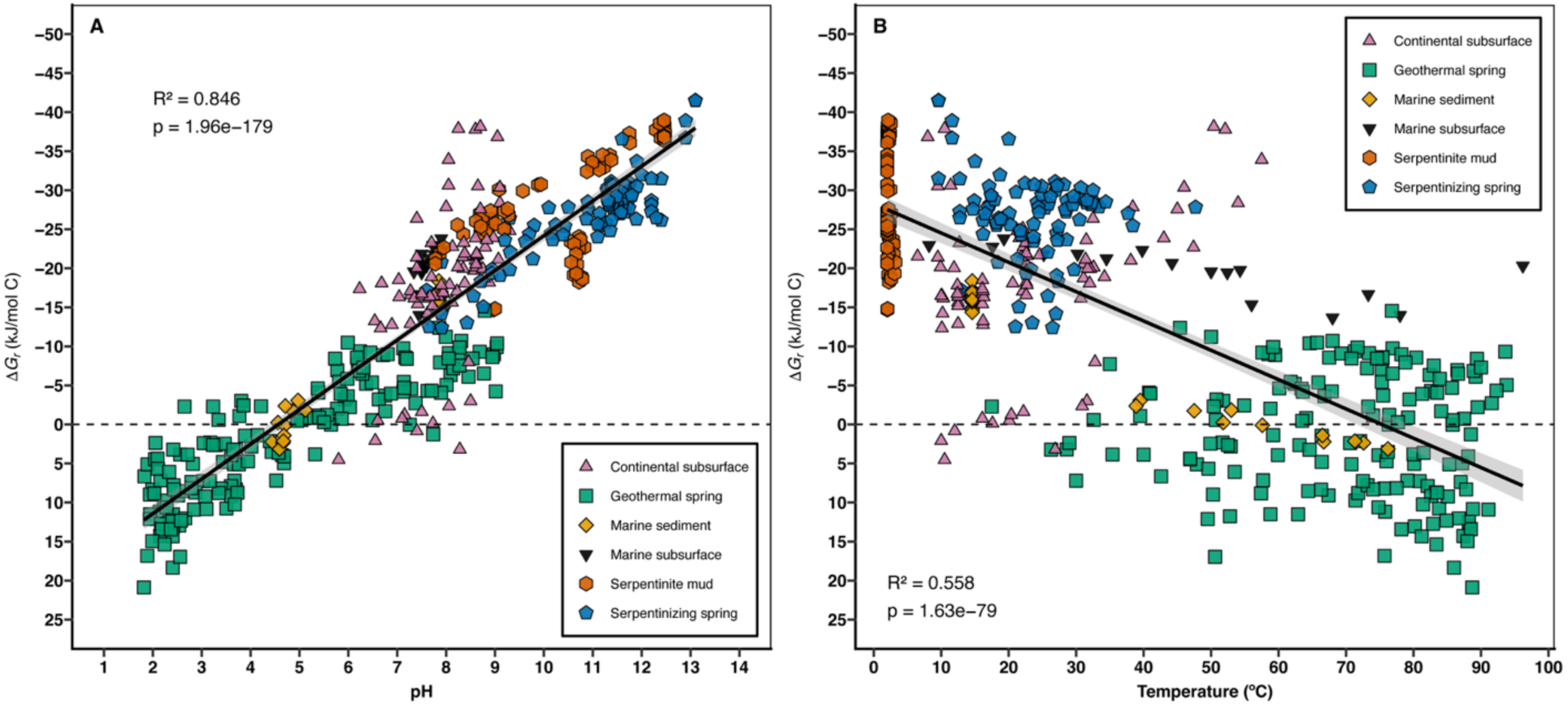
Values of the overall Gibbs energy of reaction, Δ*G_r_* (kJ/mol carbon), for Rxn. 1 (2CH_4(aq’)_ + 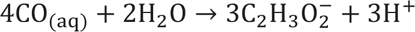) represented as a function of A) pH and B) temperature (°C) from 439 samples from natural anoxic environments. Temperature, pH, and activities of CH4, CO, and acetate were sourced from *in situ* geochemical data from six environment types, represented as the shape of the points.

### Modeled Martian and early Earth fluids

Modeled fluid composition for early Earth and Mars spanned a wide energetic range for Rxn. 1, and was exergonic across all 186 scenarios evaluated, with Δ*G_r_* values spanning from -8 to -43 kJ/mol C (Fig. 5). Under 22 modeled Martian fluid scenarios, including Eridania Basin hydrothermal fluid models (Rucker et al., 2023) and the Mars geochemical models from Marlow et al. (2014), under low-and high-CO atmospheric conditions, Δ*G_r_* ranged from -8 to –43 kJ/mol C. Early Earth ocean compositions (164 modeled scenarios) yield negative Δ*G_r_* values. The four atmospheric endmembers (CH₄ partial pressures of 5×10^-3^ or 1.8×10^-2^ bar and CO partial pressures of 10^-5^or 10^-2^ bar) produced dissolved CH4 concentrations of 4.85 to 38.5 µM and dissolved CO concentrations of 8.2 nM to 16.9 µM over a temperature range of 0 to 40 °C. Under these conditions, Δ*G_r_* spans from - 9 to -25 kJ mol C. Six representative points (high or low gas partial pressures, at 10, 25, and 40 °C), selected to span endmember conditions and an intermediate temperature, are shown in Fig. 5 to illustrate the full range of predicted Δ*G_r_*.

### Energy yields of other reactions involving CO, CH4, and acetate

Δ*G_r_* was calculated for six additional reactions (Rxns. 6 through 11) representing known metabolisms that either produce acetate or consume CO and CH4 under the geochemical conditions of the 439 natural samples (Fig. 7). Energy yields were normalized by moles of carbon to facilitate comparisons between samples. Reaction. 1 yielded Δ*G_r_* values between +21 to -41 kJ/mol C and was exergonic in 78% of samples. Reaction 6 (water-gas shift reaction) spanned +13 to -29 kJ/mol C and was exergonic in 83% of natural samples. Reaction 7 (CO oxidation coupled to sulfate reduction) was exergonic in all samples and yielded -5 to -45 kJ/mol C. Reaction 8 (methanogenic CO disproportionation) also produced negative values of Δ*G_r_* in all samples, ranging from -7 to -45 kJ/mol C. Reaction 9 (acetogenic CO disproportionation) yielded +3 to -50 kJ/mol C and was exergonic in all but two samples. Reaction 10 (methane oxidation coupled to sulfate reduction) yielded +50 to -53 kJ/mol C and was exergonic in 57% of samples. Reaction 11 (homoacetogenesis) was exergonic in 86% of samples and yielded Δ*G_r_* values between +37 and -96 kJ/mol C.

**Figure 7.**
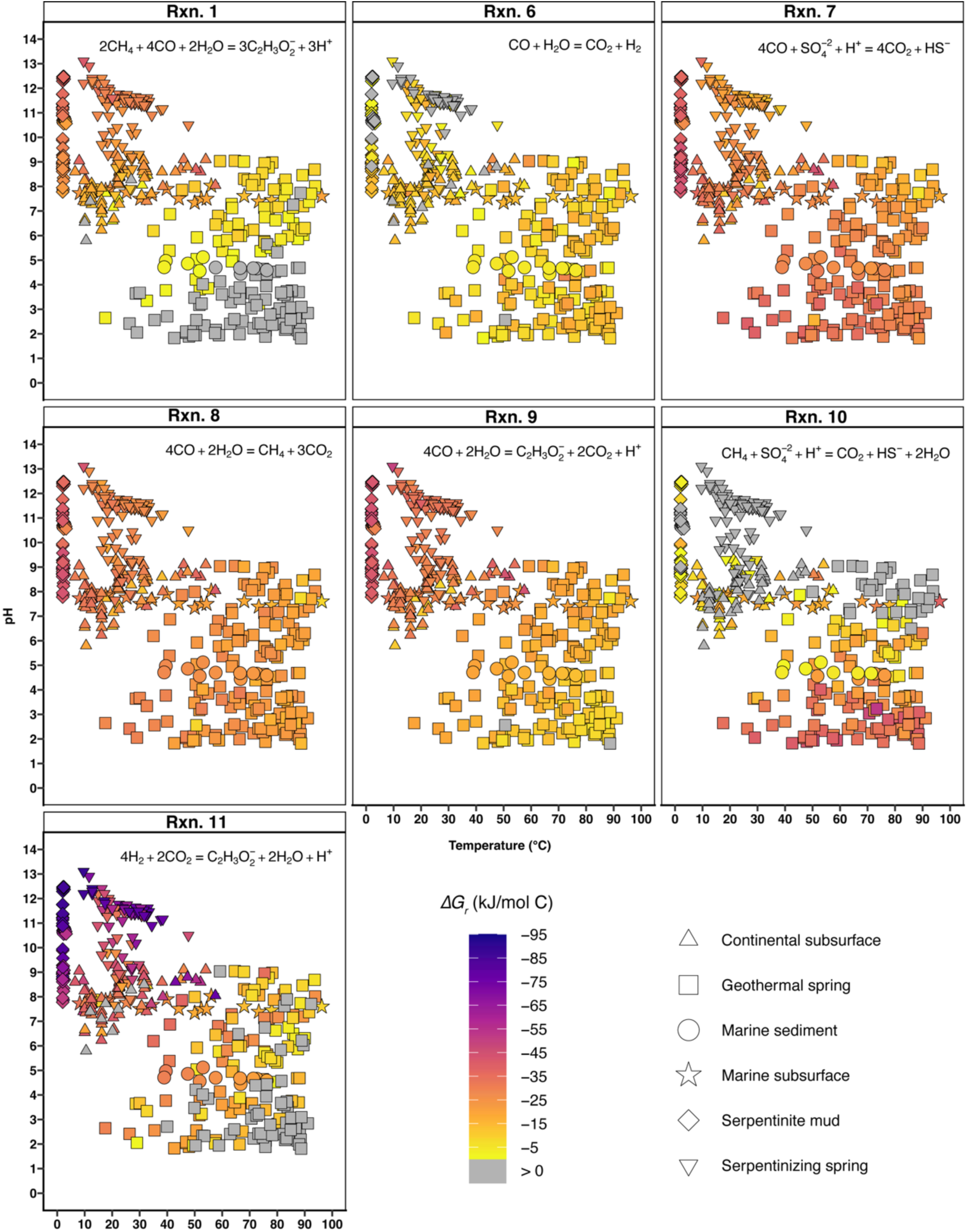
Values of the overall Gibbs energy of reaction, Δ*G_r_*, normalized per mole of carbon for Rxn. 1 and reference reactions (Rxns. 6-11). Values of Δ*G_r_* are shown as a function of temperature and pH for 467 combinations of temperature, pH, pressure, and activities of CH4, CO, and acetate representing samples from seven anoxic environment types, along with modeled early Earth ocean and Martian fluids. Temperature, pH, pressure and activities of CH4, CO, and acetate were taken directly from *in situ* geochemical data from seven environment types, represented as the shape of the points. Point fill indicates the magnitude of Δ*G_r_* (kJ/mol C); gray points denote samples where Δ*G_r_* > 0.

Energy densities (*E_r_*) were calculated from Δ*G_r_* to represent the amount of energy available in a kilogram of water, normalized to the activity of the limiting reactant (Fig. 8). Because energy densities span several orders of magnitude across reactions and environments, values corresponding to exergonic reactions (Δ*G_r_* < 0) are plotted in Fig. 8 as log10(-*E_r_*) to improve visual resolution; all numerical values reported in the text refer to untransformed values of *E_r_*. Across the natural samples, Rxn. 1 yields energy densities ranging from +2.39×10^-6^ kJ/kg H2O to -3.46×10^-2^ kJ/kg H2O. Rxn. 11 spans +3.62×10^-5^ to -3.74×10^-3^ kJ/kg H2O. Rxn. 12 ranges from -9.7×10^-12^ to -4.4×10^-5^ kJ/kg H2O. Rxn. 13 yields -9.1×10^-10^ to -2.8×10^-2^ kJ/kg H2O. Rxn. 14 had energy densities between +4.2×10^-9^ and - 3.0×10^-2^. Rxn. 15 spans from +6.35×10^-7^ to -4.85×10^-4^. Rxn. 16 yields energy densities ranging from +3.57×10^-7^ to -2.96×10^-1^, which were the largest values observed.

**Figure 8.**
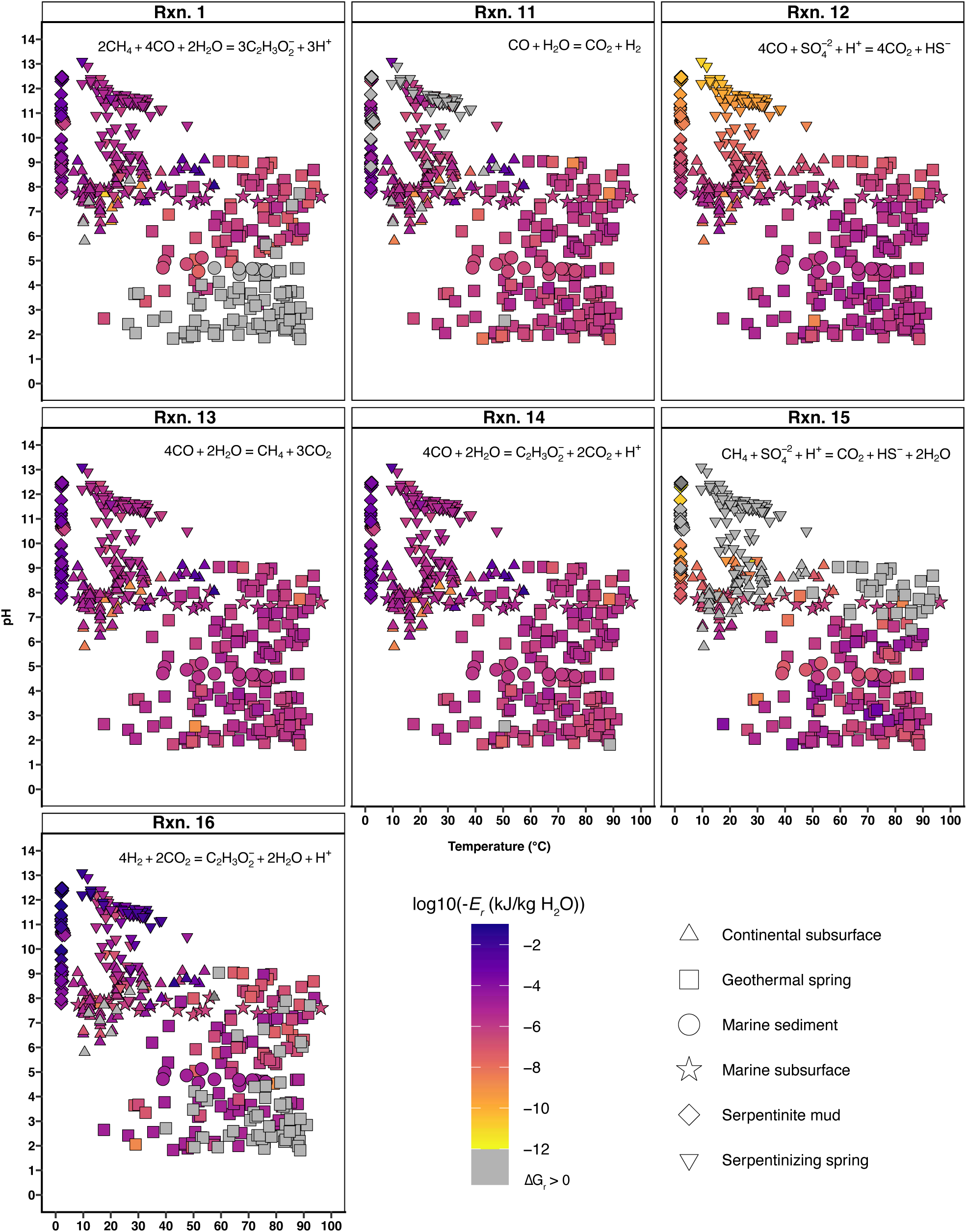
Energy densities (*E_r_*, kJ/kg H2O) for Rxn. 1 and reference reactions (Rxns. 11-16) across 439 natural samples. Energy densities represent the energy available per kilogram of water normalized by the stoichiometrically corrected activity of the limiting reactant. Temperature, pH, and activities of CH4, CO, acetate, sulfate, sulfide, bicarbonate, and H2 were sourced from *in situ* geochemical data from seven environment types, represented as the shape of the points. Point fill indicates the log10-transformed magnitude of *E_r_*, where darker colors represent more higher energy densities (more exergonic values of Δ*G_r_*); gray points denote samples where Δ*G_r_* > 0.

To evaluate relative energetic favorability across reactions, pairwise differences in the Gibbs energy and log-transformed energy density (Δ*logE*) were calculated for each sample using Reaction 1 as a reference. Comparisons based on Gibs energy differences yielded mixed outcomes across reactions, with median differences and the fraction of samples in which Rxn. 1 was more exergonic varying by reaction (Table 1). In contrast, paired comparisons based on energy density showed that Rxn. 1 yielded higher energy densities than most alternative reactions in the majority of samples in which both reactions were exergonic. Median Δ*logE* values were negative for all reactions except homoacetogenesis and differed significantly from zero for all reaction comparisons (Wilcoxon signed-rank test, Holm-corrected p < 0.001; Table 1).

**Table 1.** Paired comparisons of energy density between Reaction 1 and alternative anaerobic metabolic reactions across 439 natural samples. Differences are reported as Δ*logE* = *log_10_*(−*E_r_*) – *log*_10_(−*E_r_*), where negative values indicate higher energy density for Reaction 1. Comparisons are restricted to samples in which both reactions are exergonic. For each reaction, the table reports the number of paired samples (*n*), the median ΔlogE with interquartile range (Q25-Q75), and the fraction of samples in which Rxn.1 is more exergonic based on pairwise Gibbs energy comparisons. Reported p-values (*padj*) correspond to Wilcoxon signed-rank tests of median Δ*logE* against zero and are Holm-corrected for multiple comparisons.

| Reaction Number | Reaction | *n* | Median $\Delta\log E$ | Q25 | Q75 | Fraction of samples where Rxn.1 is more exergonic | $p_{adj}$ |
| --- | --- | --- | --- | --- | --- | --- | --- |
| 6 | Water-gas shift | 275 | -0.31 | -0.54 | 0.17 | 0.68 | $8.6 \times 10^{-6}$ |
| 7 | CO oxidation via sulfate reduction | 344 | -1.00 | -3.81 | 0.09 | 0.72 | $1.3 \times 10^{-25}$ |
| 8 | Methanogenic CO disproportionation | 344 | -0.004 | -0.17 | 0.38 | 0.51 | $5.9 \times 10^{-4}$ |
| 9 | Acetogenic CO disproportionation | 344 | -0.003 | -0.10 | 0.34 | 0.51 | $2.2 \times 10^{-4}$ |
| 10 | Methane oxidation via sulfate reduction | 163 | -1.33 | -3.63 | 0.05 | 0.75 | $4.1 \times 10^{-13}$ |
| 11 | Homoacetogenesis | 327 | +0.45 | -0.38 | 1.83 | 0.39 | $6.8 \times 10^{-10}$ |

## Discussion

Although the absolute limits of life are ultimately constrained by universal physical and chemical principles, such as the thermal stability of macromolecules and the requirements for liquid water and redox chemistry, the physicochemical boundaries within which life has been observed are largely empirically defined. The known limits of life continue to expand as new extremophilic microorganisms and previously unrecognized metabolic strategies are discovered, particularly in environments that serve as analogs for extraterrestrial settings (Jones et al., 2018; Merino et al., 2019). Each expansion of life’s observed limits broadens the range of conditions under which biology might persist elsewhere in the Universe and adds to the repertoire of biochemical and energetic strategies available to microorganisms. Because life beyond Earth would be governed by the same thermodynamic laws that constrain terrestrial systems, discoveries on Earth directly inform predictions about possible metabolisms on other planetary bodies as well as early Earth. Although many ecological and biochemical factors influence habitability (Cockell et al., 2016), energy remains a universal requirement (Hoehler, 2007; Hoehler et al., 2007; Shock & Holland, 2007). Identifying which redox processes are exergonic under specific geochemical conditions is therefore a foundational step in evaluating whether an environment could sustain microbial life.

Thermodynamics provides the universal framework for these predictions by revealing which redox reactions are energetically favorable under defined environmental conditions. This approach has been widely applied to interpret microbial processes in extreme environments (Canovas et al., 2017; Howells et al., 2025; Shock et al., 2010; Vick et al., 2010) and for constraining potential metabolisms in settings where direct observations are limited or difficult to acquire (Alain et al., 2022; Amend et al., 2011; Boettger et al., 2013; Bojanova et al., 2023; McCollom, 2000; Osburn et al., 2014). Importantly, thermodynamic analyses allow us to identify reactions that are theoretically available for microorganisms to exploit, independent of whether those reactions are known to occur in modern ecosystems. This capacity expands the set of reactions considered in discussions of habitability, early metabolic evolution, and planetary biogeochemistry where empirical measurements of life are not yet possible (Hand et al., 2007; Jakosky & Shock, 1998; Marlow et al., 2014; Rucker et al., 2023; Shock, 1997; Yanez et al., 2024; Zolotov & Shock, 2003).

Carbon comproportionation provides a useful test case for applying this thermodynamic framework to an underexplored class of redox reactions. Although comproportionation chemistry has been proposed and demonstrated for reactions involving nitrogen and sulfur (Amend et al., 2020; Aronson et al., 2025; Van de Graaf et al., 1995), analogous reactions involving carbon have received relatively little attention, despite carbon’s central role in both microbial metabolism and global biogeochemical cycles. In principle, carbon species spanning a wide range of oxidation states can participate in redox reactions that yield products of intermediate oxidation states. While this reaction space has been examined for the energetics of fermentation reactions (D. E. LaRowe & Amend, 2019), carbon comproportionation has not been examined as a distinct class of reactions in the context of microbial energy conservation. Exploring this reaction space therefore provides a thermodynamically grounded approach for identifying previously unrecognized carbon transformations that may fall within the energetic limits of life, particularly in the extreme and/or low-energy environments relevant to early Earth and planetary settings.

### A survey of carbon comproportionation reactions

The distribution of nominal oxidation states of carbon (NOSC) provides a quantitative structure for exploring carbon comproportionation reactions. The standard state Gibbs energy of carbon oxidation (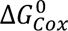) is inversely correlated with NOSC (Fig. 1) (D. E. LaRowe & Van Cappellen, 2011). Although 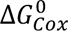 can vary by >100 kJ/mol C among compounds sharing the same NOSC, which may be attributed to differences in molecular structure including the presence of heteroatoms such as sulfur, nitrogen, and oxygen, the overall trend indicates that NOSC is a robust predictor of the energetics of carbon oxidation. This relationship is useful when evaluating carbon comproportionation reactions. Compounds with more negative NOSCs, and therefore more positive values of 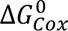, represent potential electron donors in comproportionation reactions, whereas compounds with higher NOSCs and more positive values of 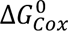 represent potential electron acceptors.

However, most combinations represented are unlikely to support microbial catabolisms. Many of the most reduced carbon compounds occur as high-molecular weight hydrocarbons that are poorly soluble, degraded slowly in natural environments and infrequently serve as direct catabolic substrates for anaerobic microorganisms (Hu et al., 2025). For this reason, the reaction set in Fig. 2 was restricted to small, labile, water-soluble compounds that are frequently involved in microbial carbon transformations. Even within this restricted set of reactions, standard state Gibbs energies alone do not identify physiologically plausible reactions. Negative values of 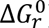 provide an initial indication of thermodynamic favorability, but reactions with positive standard states can become exergonic under certain environmental conditions. For example, elemental sulfur disproportionation exhibits a standard state Gibbs energy of +125 kJ mol^-1^ at 25 °C, yet still supports microbial growth because natural conditions can drive the overall Gibbs energy negative (Aronson et al., 2023). Accordingly, reactions with both negative and positive 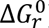 values were considered in this initial screening, with the expectation that thermodynamic favorability under natural conditions may differ from standard state values.

By applying thermodynamic, geochemical, and biochemical considerations we can further refine the set of carbon comproportionation reactions that may serve as potential microbial catabolisms. Several acetylene-based reactions in Fig. 2 yield negative 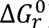 values; however, acetylene is infrequently quantified in natural anoxic environments. Although acetylene disproportionation is an established microbial metabolism (Akob et al., 2018) and provides an energetically favorable route for acetylene utilization (>70 kJ/mol) (Yanez et al., 2024), the scarcity of acetylene concentration measurements limits the ability to evaluate acetylene-based comproportionation reactions across environmentally relevant ranges of pH, temperature, and fluid composition. In contrast to CO- and CH4-based reactions, which can be assessed across hundreds of natural samples, acetylene-based pathways cannot presently be evaluated at comparable scale. Independent of substrate availability constraints, many of the remaining thermodynamically plausible carbon comproportionation reactions represent simple reversals of fermentation reactions (e.g., CO2 + ethanol → glucose). Such reversed fermentation reactions would require substantial ATP investment because forward fermentation relies on substrate-level phosphorylation to conserve energy. Reversing these pathways would therefore impose a net energetic cost and provide no mechanism for ATP generation, making them physiologically unrealistic despite favorable values of 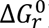.

After applying these constraints, a smaller number of reactions remain plausible as potential catabolic pathways. Among these, CH4 oxidation coupled to CO reduction to form acetate (Rxn. 1) emerged as a compelling candidate, hereafter ‘acetogenic CH4-CO comproportionation’. This reaction links two small, soluble, widely co-occurring carbon species and produces acetate, a central intermediate and common end product of anaerobic carbon metabolism, including acetogenic carbon fixation and fermentation. Acetogenic CH4-CO comproportionation spans a large redox gradient (oxidation states: +2, -4, and 0), and can be evaluated across hundreds of natural geochemical samples containing measured CO and CH4, which sets this reaction apart from many of those in Fig. 2. These attributes make acetogenic CH4-CO comproportionation an environmentally grounded case study for assessing carbon comproportionation as a potential mode of energy conservation. Moreover, these attributes enable environment-specific evaluation of both Gibbs energy and energy density, allowing this reaction to be assessed not only in theory but also within the geochemical contexts where it may realistically occur.

### Environmental controls on the energetics of acetogenic CH₄–CO comproportionation

The energetic behavior of Rxn. 1 reflects a combination of intrinsic reaction properties and strong influence of environmental chemistry. The standard state Gibbs energy (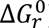) defines the baseline energetics of the reaction as a function of temperature and pressure, whereas pH and reactant activities determine the overall favorability of the reaction under specific chemical conditions. Across the temperature and pressure ranges examined, variations in 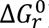 for Rxn. 1 are modest, on the order of several kilojoules per mole of carbon, indicating that pressure exerts only a minor influence on reaction energetics relative to environmental chemistry (Fig. 3). When Δ*G_r_* is evaluated at environmentally realistic activities of CH4, CO, and acetate, Rxn. 1 exhibits a strong dependence on pH (Fig. 4). This dependence arises directly from reaction stoichiometry, as the net consumption of three protons strongly favors alkaline conditions; pH therefore exerts a first-order control on Δ*G_r_*, while temperature plays a secondary role by modulating 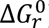. These controls produce systematic patterns across natural environments, with alkaline, reducing systems clustering in the most exergonic region of Δ*G_r_* space (Figs. 5 and 6A).

Among the environments examined, continental and marine serpentinizing systems consistently support the most favorable energetics for Rxn. 1. These systems combine high pH, low to moderate temperatures, and highly reducing conditions that produce both CH4 and CO. In serpentinizing systems, CH4 is formed by CO2 reduction with the abundant H2 that is produced through the serpentinization reaction (Horita & Berndt, 1999; Leong & Shock, 2020; McCollom & Seewald, 2013). CO can also form abiotically in these systems through the reduction of CO2 to form formic acid, followed by the decomposition of formic acid to CO (McCollom & Seewald, 2007; Seewald et al., 2005). In these settings, dissolved inorganic carbon is often sequestered as carbonate minerals leaving fluids depleted in bioavailable CO2/HCO3-for autotrophic carbon fixation. Under such conditions, carbon species such as formate or CO may serve as important carbon sources (McCollom & Seewald, 2013), and, in the absence of oxidants such as oxygen, sulfate, nitrate, or ferric iron, may function as terminal electron acceptors.

Deep continental subsurface fluids frequently support favorable energetics for Rxn. 1 across a broad range of conditions, but exhibit some of the greatest variability in energy yields. These systems commonly maintain circumneutral to alkaline pH, low oxidant availability, and long-term isolation from surficial inputs, yet differ significantly in the availability of CH4 and CO, which can be derived from abiotic reactions and thermogenic processes (G. Holland et al., 2013; Kieft et al., 2005; Sherwood Lollar et al., 1993, 2006). As a result, continental subsurface environments host some of the highest and lowest energy yields calculated for Rxn. 1. Sites within the Witwatersrand Basin, for example, exhibit exceptionally high CO concentrations (680 µM at Kloof Mine) and support some of the most favorable energetics, whereas fluids from the Deep Mine Microbial Observatory (DeMMO) contain extremely low (sub nanomolar) concentrations of both CO and CH4 and yield correspondingly low or endergonic values of Δ*G_r_*. This contrast highlights that within otherwise similar subsurface geochemical regimes, substrate availability controls the energy yield of Rxn. 1.

In contrast, geothermal springs and hydrothermal sediments exhibit highly variable energetics, which reflect their geochemical heterogeneity. Acidic, high-temperature systems tend to yield less favorable (more endergonic) values of Δ*G_r_*, consistent with the dependence of Rxn. 1 on high pH and lower temperature. Geothermal springs span some of the widest ranges of pH and temperature on Earth, from strongly acidic to alkaline fluids (pH 1.8 to 9.0) and from moderate to near-boiling temperatures (17.5 to 93.8 °C), resulting in large ranges in predicted energy yields. Alkaline geothermal springs, however, yield favorable energetics despite elevated temperatures, again highlighting the dominant role of pH over temperature in controlling Δ*G_r_* (Fig. 6). Marine sediments likewise exhibit broad variability in the energetics of Rxn. 1. In hydrothermally influenced settings such as Guaymas Basin, thermolytic cracking of organic matter and magmatic inputs generate both CH4 and CO (Conrad & Seiler, 1985; Lizarralde et al., 2023; Von Damm et al., 1985), while microbial methanogenesis contributes substantially to the CH4 pool, which can exceed 100 mM at *in situ* pressures and temperatures (Bojanova et al., 2023). Despite high temperatures, these fluids support exergonic conditions for Rxn. 1 due to the high activities of reactants. In contrast, the hot and acidic shallow-sea hydrothermal sediments from Milos, Greece yield the least favorable estimates of Δ*G_r_* for Rxn. 1, due to the combined low pH and high temperature (Figs. 3, 4, 6).

Across all environments studied here, the most favorable conditions for Rxn. 1 occur where high pH and persistent sources of both reactants co-occur. These patterns indicate that the energetic feasibility of Rxn. 1 is largely constrained by the geochemical and geological contexts that sustain stable fluxes of CH4 and CO over timescales sufficient for microbial utilization. Such environments provide the necessary foundation for evaluating whether and where Rxn. 1 is a viable microbial catabolism.

### Comparison of acetogenic CH₄–CO comproportionation with known metabolic reactions

The Gibbs energy of Rxn. 1 was compared directly to six established anaerobic catabolisms involving CO, CH₄, and acetate (Rxns. 11–16) across all 439 natural samples to place acetogenic CH₄–CO comproportionation within the thermodynamic context of known microbial metabolisms (Fig. 7). When normalized per mole of carbon, Rxn. 1 spans a similar energetic range to these reactions and is frequently more exergonic than under the same environmental conditions. However, comparisons based on Δ*G_r_* alone indicate that Rxn. 1 is not consistently the most exergonic reaction available under shared environmental conditions, particularly relative to CO disproportionation (Rxns. 8 and 9), CO oxidation coupled to sulfate reduction (Rxn. 10), and homoacetogenesis (Rxn. 11). This highlights the limitations of evaluating energetic competitiveness solely on the basis of Gibbs energy yields.

Although Gibbs energies constrain the theoretical energy yield of a reaction, they do not reflect whether the required substrates are present at activities that allow that energy to be accessed by microorganisms. Expressing Gibbs energies solely on a per-mole or per-electron basis can therefore be misleading, particularly when substrates are present at extremely low concentrations. Normalizing Gibbs energies to the availability of the limiting reactant (i.e., expressing them as energy densities per unit mass or volume), provides a more ecologically relevant metric that better reflects which reactions can sustain microbial populations and shape biogeochemical processes in natural environments (D. LaRowe & Amend, 2014, 2019). To evaluate the amount of energy that accessible in a given environment, energy densities (*E_r_*, kJ/kg H2O) were calculated for each reaction by weighting Gibbs energy yields by the availability of the limiting reactant (Fig. 8). For Rxn. 1, reaction stoichiometry dictates that two moles of CH4 are required for every four moles of CO, such that the energy availability depends on the relative abundances of these substrates. Across the natural samples examined here, CH4:CO activity ratios span more than eight orders of magnitude, causing Rxn. 1 to be either CO-limited or CH4-limited depending on the environment. This constraint limits the amount of energy that can be accessed even when Gibbs energies are favorable. Energy density calculations therefore reveal that the highest realizable energy yields for Rxn. 1 occur in deep continental subsurface fluids and marine and continental serpentinizing systems, whereas environments with low CO or CH4 activities yield orders-of-magnitude lower energy densities. These patterns demonstrate that the availability of limiting substrates, rather than thermodynamic favorability alone, governs the accessible energy yield of Rxn. 1 across all systems.

To compare reactions based on accessible energy, paired differences in energy density were evaluated using the log-transformed metric Δ*logE*, calculated only for samples in which both reactions were exergonic. When evaluated in this way, Rxn. 1 is energetically competitive with multiple established anaerobic catabolisms, but its relative favorability depends strongly on environmental context. Rxn. 1 exhibits higher energy densities than the water-gas shift reaction (Rxn. 6) in most samples (Table 1). Bacteria and archaea capable of catalyzing the water-gas shift reaction are typically thermophiles, with cultured representatives from anaerobic bioreactors (Alves et al., 2013; Parshina et al., 2010), hot springs (Balk et al., 2009; Novikov et al., 2011; Slepova et al., 2006, 2009; Tatyana G. Sokolova et al., 2004; Svetlichny et al., 1991), and marine hydrothermal vents (Bae et al., 2006; T G Sokolova et al., 2001) – environments in which Rxn. 1 tends to be least favorable. The energy densities of Rxn. 1 also exceed CO oxidation coupled to sulfate reduction (Rxn. 7) in many high-pH, low temperature environments. In sulfate-reducers, CO oxidation typically proceeds via the water-gas shift reaction (Rxn. 6) with CO first converted to H2 and CO2, and the resulting H2 powering sulfate reduction (Oelgeschläger & Rother, 2008; Parshina et al., 2005, 2010). This has been interpreted as a strategy to detoxify CO, as many sulfate reducers are inhibited by CO (Parshina et al., 2010). In many of the hyperalkaline sites (c.f. terrestrial serpentinizing) proton and/or sulfate limit the accessible energy yield of Rxn. 7, while Rxn. 1 remains energetically favorable.

In contrast, Rxn. 1 and CO disproportionation Rxns. 8 and 9, yield similar quantities of energy, as reflected by near-zero median Δ*logE* values. This overlap indicates direct competition for CO between Rxn. 1 and 8 or 9 under many environmental conditions. While CO disproportionation has been demonstrated in both methanogenic and acetogenic organisms, growth is often slow or inhibited at elevated CO concentrations (Daniels et al., 1977; Henstra & Stams, 2011; O’Brien et al., 1984; Rother & Metcalf, 2004). Methanogens capable of CO metabolism (c.f. *Methanobacterium thermoautotrophicum* and *Methanosarcina barkeri*) typically utilize Rxn. 11, with product H2 used to fuel CO2 reduction to methane (Daniels et al., 1977; O’Brien et al., 1984). In contrast, *Methanosarcina acetivorans* C2A lacks the ability to grow on H2/CO2, and instead sends CO-derived electrons into the Wood-Ljungdahl pathway to form acetate (Rother & Metcalf, 2004). Even where CO disproportionation is thermodynamically feasible, the underlying metabolic strategies and physiological constraints differ between organisms. The abundance, distribution, and ecological relevance of alternative CO-based metabolic strategies have been demonstrated in few pure cultures to-date, and remain poorly constrained. Further investigation in cultivars and in natural environments is needed to determine the ubiquity and importance of this process in nature.

The anaerobic oxidation of methane (AOM) coupled to sulfate reduction (Rxn. 15) is an obligate syntrophic metabolism common in sulfate-rich marine sediments (Knittel & Boetius, 2009). AOM with sulfate yields lower energy densities than Rxn. 1 in most exergonic sites herein, and is largely unfavorable in alkaline, sulfate-poor environments where Rxn. 1 has the highest energy densities (Fig. 8). Homoacetogenesis (Rxn. 16) frequently yields higher energy densities than Rxn. 1, but does not compete directly for substrates, such that acetogenic CH4-CO comproportionation and homoacetogenesis could coexist within the same environments. These comparisons show that when evaluated in terms of energy density rather than Gibbs energy alone, Rxn. 1 is energetically competitive with multiple known anaerobic catabolisms across a wide range of natural geochemical conditions. Accounting explicitly for substrate availability and reaction stoichiometry provides a more realistic framework for assessing the energetic viability of carbon comproportionation reactions in nature.

### Modeled fluids from Mars and early Earth

Modeled Martian and early Earth fluids allow the energetics of Rxn. 1 to be constrained under inaccesible natural conditions using independently derived atmospheric and water-rock interaction models, in an approach analogous to that applied to natural terrestrial samples. Across all 186 modeled scenarios, including 164 early Earth ocean compositions and 22 Martian hydrothermal fluid models, Δ*G_r_* values are negative, spanning -8 to -43 kJ/mol C.

Modeled Martian fluid compositions support the energetic feasibility of Rxn. 1 across a wide range of plausible subsurface conditions. Using fluid chemistries derived from Marlow et al. (2014) and Rucker et al. (2023), which represent acidic sulfate alteration fluids, alkaline basalt alteration fluids, and serpentinization-associated analogs, Rxn. 1 remains exergonic across all modeled scenarios evaluated. These fluids encompass a broad range of pH (0.67 to 11.86) and temperature (9 °C to 83 °C), reflecting both the diversity of aqueous environments proposed for ancient Mars and the uncertainty inherent in reconstructing Martian subsurface geochemistry. For each modeled fluid composition, aqueous CO concentrations were estimated from atmospheric mixing ratios of 1% and 50% using Henry’s law (Koyama et al., 2024), while aqueous CH4 activities were fixed at values adopted in the original fluid models (Marlow et al., 2014; Rucker et al., 2023). Across both low and high CO endmembers, predicted Gibbs energies overlap with those calculated for natural terrestrial subsurface environments (Fig. 5). Although estimates of Martian geochemistry necessarily span a broad range of modeled fluid compositions – many of which have already been invoked to support other anaerobic methane metabolisms – Rxn. 1 consistently falls within the thermodynamic range known to support microbial energy conservation.

For the early Earth, widely accepted models of atmospheric redox state assume a steady-state composition determined by mantle outgassing, in which CO2 is the dominant gas, with only minor amounts of CO and virtually no CH4 (Abelson, 1966; H. D. Holland, 1962, 1984; Poole, 1951). Multiple lines of theoretical and geochemical evidence, however, indicate that this steady-state framework does not capture the full range of Archean conditions. Large impacts, particularly during the late stages of accretion, are expected to have generated transient, and potentially longer-lived reducing atmospheres enriched in CO and CH4 (Hashimoto et al., 2007; Schaefer & Fegley, 2007, 2010; Urey, 1952; Zahnle et al., 2010). An alternative interpretation holds that large impacts may not consistently generate strongly reducing atmospheres, as a substantial fraction of the reducing capacity of impactors could be sequestered in the crust and upper mantle rather than directly influencing surface waters and atmospheric chemistry (Citron & Stewart, 2022).

Regardless, the ranges explored here are intended to capture plausible post-impact and mantle-buffered endmembers during intervals following the last ocean-vaporizing impacts, when long-lived surface oceans and subsurface circulation were present and capable of sustaining redox chemistry on biologically relevant timescales. Depending on impactor size and composition, reducing conditions could persist for thousands to millions of years until hydrogen escape to space restored mantle-buffered redox conditions (Wogan et al., 2023; Zahnle et al., 2010). Photochemical and impact-driven models suggest that CH4 concentrations of 5×10^-3^ to 1.8×10^-2^ bar are plausible during portions of the Archean, consistent with constraints from xenon isotope fractionation and inferred microbial methane fluxes (Catling et al., 2001; Catling & Zahnle, 2020; Kharecha et al., 2005; Ozaki et al., 2018; Wogan et al., 2023). CO abundances are less well constrained, but episodic enrichment to 10^-5^ to 10^-2^ bar has been predicted during impact-driven reducing intervals, particularly when cometary CO ice or reduced carbon is delivered and oxidized (Catling & Zahnle, 2020; Schaefer & Fegley, 2007; Wogan et al., 2023).

The early Earth ocean models therefore span both steady-state, mantle-buffered atmospheres with low CO and CH4 and transient reducing periods in which these gases are substantially enriched. That Rxn. 1 remains exergonic across this entire range indicates that its energetic feasibility does not depend on transient or extreme atmospheric conditions. Moreover, if CO and CH4 produced during transient atmospheric episodes dissolve into ocean waters and circulate at depth, their residence times may greatly exceed those of the atmosphere, particularly in subsurface environments isolated from rapid reoxidation. The decoupling of atmospheric composition and subsurface redox chemistry is consistent with the favorable energetics of Rxn. 1 observed in modern serpentinizing systems and deep subsurface fluids.

The highest energy densities for Rxn. 1 occur in deep continental subsurface fluids and serpentinizing systems, which are characterized by isolation from phototrophic energy sources, low energy fluxes, and redox disequilibria sustained by water-rock reactions. These conditions define a class of energy-limited subsurface environments that are increasingly central to astrobiological investigation. In this context, Earth’s subsurface environments provide relevant analogs for extraterrestrial environments because of shared energetic and geochemical constraints. Serpentinization represents a key process analog linking these systems. Serpentinization-hosted alkaline hydrothermal systems have also been widely invoked as plausible settings for early metabolism on Earth (Martin et al., 2008; M. J. Russell et al., 2010), particularly in models that place acetogenesis and the Wood-Ljungdahl pathway at the core of some of the earliest energy-conserving reactions (Michael J. Russell & Martin, 2004; Weiss et al., 2016). Within this framework, reactions coupling CO and CH4 to acetate formation analogous to Rxn. 1, have been discussed as plausible components of early acetogenic metabolism at alkaline hydrothermal vents (Nitschke & Russell, 2013; Michael J. Russell & Nitschke, 2017). Consistent with this view, deeply branching acetogenic lineages have been identified in serpentinite-associated subsurface fracture waters, including members of the candidate phyla *Acetothermia*, *Lithacetigenota*, and NPL-UPL2, as well as the acetogenic archaeon within the order *Methanocellales* (Colman et al., 2022; Nobu et al., 2023; Suzuki et al., 2018, 2024). While these organisms do not utilize direct CO-CH4 reactions, they employ canonical or modified Wood-Ljungdahl pathways, in some cases using CO, formate, or glycine as substrates for acetogenesis. These observations highlight the long-standing association between serpentinization-driven environments, C1 chemistry, and acetogenic energy metabolism.

Exploration targets for life detection – including Mars, Europa, and Enceladus – possess surface environments that are largely inhospitable to life, but are thought to host, or to have hosted, subsurface aqueous systems where chemical energy would govern habitability (Jones et al., 2018; Michalski et al., 2013, 2018). Evidence for serpentinization has emerged on other planetary bodies. Detection of H2 in the plume of Enceladus (Waite et al., 2017) has been attributed to ongoing water-rock serpentinization reactions in an alkaline subsurface ocean (Glein et al., 2015; Glein & Truong, 2025), while models of Europa’s ocean-seafloor interface likewise predict water-rock reactions capable of generating redox disequilibria, even where fluid pH remains poorly constrained(Hand et al., 2007; McCollom, 1999; S. Vance et al., 2007; S. D. Vance et al., 2016; S. D. Vance & Melwani Daswani, 2020; Zolotov & Shock, 2003, 2004). On Mars, the occurrence of olivine and serpentine minerals in Noachian-aged crust indicate past aqueous alteration consistent with subsurface serpentinization (Ehlmann et al., 2010; Hoefen et al., 2003; Ody et al., 2012). Taken together, these observations suggest that the geochemical conditions that favor Rxn. 1 may be present in the subsurface of other planetary bodies.

### Conclusion

In this study we identify carbon comproportionation as a previously underexplored class of anaerobic redox reactions with the potential to support microbial energy conservation in low-energy environments. We show that acetogenic CH4-CO comproportionation is exergonic across a broad range of temperatures, pH conditions, and natural fluid compositions, and furthermore that energy yield depends strongly on substrate availability rather than the Gibbs energy alone. By explicitly incorporating the availabilities of limiting reactants in each environment through energy density calculations, we demonstrate that this reaction occupies an energetic range comparable, and in some settings, exceeding, that of established anaerobic metabolisms involving CH4, CO, and acetate. The highest energy densities occur in subsurface and serpentinization-associated environments, reflecting the role of water-rock reactions in producing and maintaining CH4 and CO under reducing and often alkaline conditions. These results collectively expand the known range of energetically viable anaerobic carbon transformations, providing a quantitative framework for evaluating carbon-based catabolic reactions under realistic geochemical constraints, and motivate further investigation of carbon comproportionation reactions as potential contributors to subsurface biogeochemical cycling on Earth, and by extension, on other planetary bodies.

## Acknowledgements

H.S.A. is supported by the Energy & Biosciences-Shell Postdoctoral Fellowship. D.E.L. acknowledges financial support from the NASA Habitable Worlds program under grant 80NSSC20K0228, the NASA Exobiology program under grant NNH22ZDA001N, Southern Methodist University and the Claude C. Albritton, Jr. Endowment. W.D.L. acknowledges support from the University of Utah and NASA Exobiology program under grant 80NSSC23K1353 and thanks Alex Bradley for early conversations about on carbon dioxide reduction powered by microbial methane oxidation.

## Author contributions

W.D.L. and H.S.A. designed the study. All authors collected data from published studies. H.S.A. performed data analyses. H.S.A. wrote the manuscript with input from all co-authors. All authors reviewed and edited the manuscript. The authors declare there are no conflicts of interest for this manuscript.

## Data availability statement

All data are available in the supplementary materials. Data and R scripts written for data analysis and visualization are also hosted at https://doi.org/10.5281/zenodo.18158636.

## Supplementary Methods

Water pressure at seafloor was calculated for marine subsurface sediment samples using

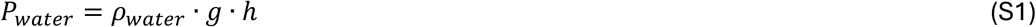

where ρ*_water_* corresponds to density (1035 kg/m^3^), *g* is gravity, and *h* is water depth (m).

Sediment porosity at depth z meters below seafloor, *φ_Z_*, was calculated following (Bradley et al., 2019) using

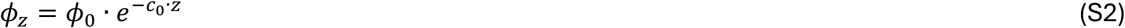

where porosity at the sediment-water interface (φ_0_) is equal to 0.87 and compaction length scale (*c*_0_) is equal to 0.00085 per meter (Bradley et al., 2019). Wet bulk density (*ρ_sed_*) of sediments was calculated using

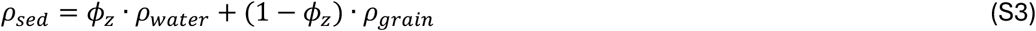

where grain density, *ρ_grain_*, is equal to 2300 kg/m^3^ (Bradley 2019). Pressure at depth z below seafloor (*P_Z_*) was calculated with

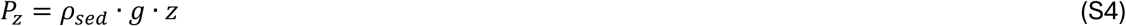

and total pressure, *P_T_*, was calculated with

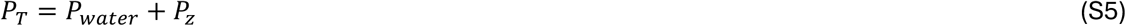

Thermodynamic data used in this study are implemented in the OrganoBioGeoTherm database distributed with CHNOSZ (Jeffrey M. Dick, 2019). This database aggregates thermodynamic properties for organic and inorganic aqueous species from the primary literature (Accornero et al., 2010; Akilan et al., 2006; N. N. Akinfiev et al., 2006; N. N. Akinfiev & Tagirov, 2014; N. N. Akinfiev & Zotov, 2010, 2023; Nikolay N. Akinfiev et al., 2020; Nikolay N. Akinfiev & Diamond, 2003; Nikolay N. Akinfiev & Plyasunov, 2014; N.N. Akinfiev & Zotov, 2001; J P Amend & Shock, 2001; Jan P. Amend & Helgeson, 1997; Jan P. Amend & Plyasunov, 2001; Azadi et al., 2019; Bandura & Lvov, 2005; Barin & Knacke, 1973; Bazarkina et al., 2010; Berman, 1988, 1990; Bird et al., 2011; Bowers & Helgeson, 1983; Boyer & Shock, 2023; Canovas & Shock, 2016; Dale et al., 1997; Denbigh, 1981; I. Diakonov et al., 1996; I. I. Diakonov et al., 1998; J. M. Dick et al., 2006; Jeffrey M. Dick, 2019; Jeffrey M. Dick et al., 2013; Jeffrey Michael Dick, 2007; Evans, 1990; Facq et al., 2014; GEOPIG, 2019; A. B. Goldberg et al., 1981; R. N. Goldberg et al., 2002; Gottschalk, 2004; Grenthe et al., 2021; Grevel & Majzlan, 2009; Gysi et al., 2016; Haar et al., 1984; Haas et al., 1995; Haas & Shock, 1999; Hakin et al., 1994; Hawrylak et al., 2006; Helgeson, 1985; Helgeson et al., 1978, 1998, 2009; B. S. Hemingway, 1982; Bruce S. Hemingway et al., 1991; Ho & Palmer, 1997; Hu et al., 2025; Huang & Sverjensky, 2019; Huston & Bastrakov, 2024; Jackson & Helgeson, 1985; Jacob et al., 2009; Johnson et al., 1992a, 1992b; Jørgensen et al., 2019; Kelley, 1960; Kitadai, 2014, 2015; Knight et al., 2024; R.J.M. Konings & Kovács, 2003; Rudy J. M. Konings et al., 2014; Langmuir et al., 2006; D. LaRowe & Amend, 2019; D. E. LaRowe & Amend, 2016, 2019; D. E. LaRowe & Dick, 2012; D. E. LaRowe & Helgeson, 2006b, 2006a; Laskar et al., 2025; Lemke et al., 2009; Leong & Shock, 2020; W. Liu et al., 2006, 2011; X. Liu et al., 2021, 2023; X. Liu & Xiao, 2020; Loges et al., 2013; Lowe et al., 2017; J. Majzlan et al., 2004; Juraj Majzlan et al., 2006; Juraj Majzlan, Grevel, et al., 2003; Juraj Majzlan, Lang, et al., 2003; Málek et al., 2011; Marini & Accornero, 2007, 2010; Martín & Delgado Soleri Gil, 2010; McCollom & Shock, 1997; Mercury et al., 2001; A. Migdisov et al., 2016, 2025; Art. A. Migdisov et al., 2009; Miron et al., 2016, 2017; Morss & Konings, 2004; Murphy & Shock, 1999; Naumov et al., 1974; Navrotsky et al., 2015; Nordstrom & Archer, 2003; Oelkers & Helgeson, 1990; O’Hare & Curtiss, 1995; Pankratz, 1970; Pankratz et al., 1987; Pankratz & King, 1970; Pankratz & Mrazek, 1982; Parker & Khodakovskii, 1995; Perfetti et al., 2008; Plummer & Busenberg, 1982; Plyasunov & Shock, 2001; Pokrovski et al., 2014; Pokrovski & Dubessy, 2015; Pokrovski & Schott, 1998; Pokrovskii & Helgeson, 1995, 1997; Popa & Konings, 2006; Prapaipong et al., 1999; Reardon & Armstrong, 1987; Richard, 2001; Richard & Gaona, 2011; Richard & Helgeson, 1998; Robie et al., 1979; Robie & Hemingway, 1972, 1995; Robinson et al., 2024; Ruaya & Seward, 1987; Sassani & Shock, 1998; M. Schulte, 2010; M. D. Schulte et al., 2001; M. D. Schulte & Rogers, 2004; M. D. Schulte & Shock, 1993; Senoh et al., 1998; Sharygin et al., 2006; Everetr L. Shock & Koretsky, 1995; Everett L Shock, 1992; Everett L. Shock, 1993, 1995; Everett L Shock et al., 1989; Everett L. Shock, Sassani, & Betz, 1997; Everett L. Shock, Sassani, Willis, et al., 1997; Everett L. Shock & Helgeson, 1988, 1990; Everett L. Shock & Koretsky, 1993; Everett L. Shock & McKinnon, 1993; Y. V. Shvarov & Bastrakov, 1999; Yu. V. Shvarov, 2008, p. 200; Shvedov & Tremaine, 1997; St Clair et al., 2019, p. 2; Stefánsson, 2001; Stefánsson et al., 2013, 2014; Stoffregen et al., 2000; D. A. Sverjensky et al., 1997; Dimitri A. Sverjensky et al., 1991, 2014, 2014; B. R. Tagirov et al., 1997, 2015, 2024; B. Tagirov & Schott, 2001; Boris R. Tagirov et al., 2013, 2025; Tardy et al., 1997; Trofimov et al., 2023; O. Vidal et al., 1992; Olivier Vidal et al., 2001, 2005; Von Der Heyden et al., 2024; Wagman et al., 1982; Williams-Jones & Vasyukova, 2022; Wood & Samson, 2000; C. Zhu & Sverjensky, 1992; Y. Zhu et al., 2005; Ziemer & Woolley, 2007; Zimmer et al., 2016).

## References

Abelson, P. H. (1966). Chemical events on the primative Earth. Proceedings of the National Academy of Sciences, 55(6), 1365–1372. 10.1073/pnas.55.6.1365

Akob, D. M., Sutton, J. M., Fierst, J. L., Haase, K. B., Baesman, S., Luther, G. W., et al. (2018). Acetylenotrophy: a hidden but ubiquitous microbial metabolism? FEMS Microbiology Ecology, 94(8), fiy103. 10.1093/femsec/fiy103

Alain, K., Aronson, H. S., Allioux, M., Yvenou, S., & Amend, J. P. (2022). Sulfur disproportionation is exergonic in the vicinity of marine hydrothermal vents. Environmental Microbiology, 24(5), 2210–2219. 10.1111/1462-2920.15975

Alves, J. I., van Gelder, A. H., Alves, M. M., Sousa, D. Z., & Plugge, C. M. (2013). Moorella stamsii sp. nov., a new anaerobic thermophilic hydrogenogenic carboxydotroph isolated from digester sludge. International Journal of Systematic and Evolutionary Microbiology, 63(Pt_11), 4072–4076. 10.1099/ijs.0.050369-0

Amend, J. P., McCollom, T. M., Hentscher, M., & Bach, W. (2011). Catabolic and anabolic energy for chemolithoautotrophs in deep-sea hydrothermal systems hosted in different rock types. Geochimica et Cosmochimica Acta, 75(19), 5736–5748. 10.1016/j.gca.2011.07.041

Amend, J. P., Aronson, H. S., LaRowe, D. E., & Macalady, J. L. (2019). Catabolisms ‘Missing in Nature.’ In 2019 Astrobiology Science Conference. AGU. Retrieved from https://agu.confex.com/agu/abscicon19/mediafile/ExtendedAbstract/Paper481351/2019-AbSciCon%20%28Amend%20et%20al._a%29_v2.pdf

Amend, J. P., Aronson, H. S., Macalady, J., & LaRowe, D. E. (2020). Another chemolithotrophic metabolism missing in nature: sulfur comproportionation. Environmental Microbiology, 22(6), 1971–1976. 10.1111/1462-2920.14982

Aronson, H. S., Clark, C. E., LaRowe, D. E., Amend, J. P., Polerecky, L., & Macalady, J. L. (2023). Sulfur disproportionating microbial communities in a dynamic, microoxic-sulfidic karst system. Geobiology, 00, 1–13. 10.1111/gbi.12574

Aronson, H. S., LaRowe, D. E., Macalady, J. L., & Amend, J. P. (2025). Isolation of a putative sulfur comproportionating microorganism. Scientific Reports, 15(1), 17999. 10.1038/s41598-025-01009-y

Bae, S.-S., Kim, Y.-J., Yang, S.-H., Lim, J.-K., Jeon, J.-H., Lee, H.-S., et al. (2006). Thermococcus onnurineus sp. nov., a Hyperthermophilic Archaeon Isolated from a Deep-Sea Hydrothermal Vent Area at the PACMANUS Field. Journal of Microbiology and Biotechnology, 16(11), 1826–1831.

Balk, M., Heilig, H. G. H. J., van Eekert, M. H. A., Stams, A. J. M., Rijpstra, I. C., Sinninghe-Damsté, J. S., et al. (2009). Isolation and characterization of a new CO-utilizing strain, Thermoanaerobacter thermohydrosulfuricus subsp. carboxydovorans, isolated from a geothermal spring in Turkey. Extremophiles, 13(6), 885–894. 10.1007/s00792-009-0276-9

Barnes, R. O., & Goldberg, E. D. (1976). Methane production and consumption in anoxic marine sediments. Geology, 4(5), 297–300. 10.1130/0091-7613(1976)4%253C297:MPACIA%253E2.0.CO;2

Boetius, A., Ravenschlag, K., Schubert, C. J., Rickert, D., Widdel, F., Gleseke, A., et al. (2000). A marine microbial consortium apparently mediating anaerobic oxidation methane. Nature, 407(6804), 623–626. 10.1038/35036572

Boettger, J., Lin, H.-T., Cowen, J. P., Hentscher, M., & Amend, J. P. (2013). Energy yields from chemolithotrophic metabolisms in igneous basement of the Juan de Fuca ridge flank system. Chemical Geology, 337*–*338, 11–19. 10.1016/j.chemgeo.2012.10.053

Bojanova, D. P., De Anda, V. Y., Haghnegahdar, M. A., Teske, A. P., Ash, J. L., Young, E. D., et al. (2023). Well-hidden methanogenesis in deep, organic-rich sediments of Guaymas Basin. The ISME Journal, 17(11), 1828–1838. 10.1038/s41396-023-01485-y

Boulart, C., Chavagnac, V., Monnin, C., Delacour, A., Ceuleneer, G., Hoareau, G., et al. (2013). differences in gas venting from ultramafic-hosted warm springs: the example of Oman and Voltri Ophiolites. Ofioliti, 38(2), 143–156. 10.4454/ofioliti.v38i2.423

Boyer, G., Robare, J., Nuri, P., Ely, T., & Shock, E. (2024). AqEquil: Python package for aqueous chemical speciation (Version 0.18.0) [Zenodo]. Retrieved from 10.5281/zenodo.10476850

Broda, E. (1977). Two kinds of lithotrophs missing in nature. Zeitschrift Für Allgemeine Mikrobiologie, 17(6), 491–493. 10.1002/jobm.19770170611

Canovas, P. A., Hoehler, T., & Shock, E. L. (2017). Geochemical bioenergetics during low-temperature serpentinization: An example from the Samail ophiolite, Sultanate of Oman. Journal of Geophysical Research: Biogeosciences, 122(7), 1821–1847. 10.1002/2017JG003825

Cardace, D., Meyer-Dombard, D. R., Woycheese, K. M., & Arcilla, C. A. (2015). Feasible metabolisms in high pH springs of the Philippines. Frontiers in Microbiology, 6. 10.3389/fmicb.2015.00010

Catling, D. C., & Zahnle, K. J. (2020). The Archean atmosphere. Science Advances, 6(9), eaax1420. 10.1126/sciadv.aax1420

Catling, D. C., Zahnle, K. J., & McKay, C. P. (2001). Biogenic Methane, Hydrogen Escape, and the Irreversible Oxidation of Early Earth. Science, 293(5531), 839–843. 10.1126/science.1061976

Citron, R. I., & Stewart, S. T. (2022). Large Impacts onto the Early Earth: Planetary Sterilization and Iron Delivery. The Planetary Science Journal, 3(5), 116. 10.3847/PSJ/ac66e8

Cockell, C. S., Bush, T., Bryce, C., Direito, S., Fox-Powell, M., Harrison, J. P., et al. (2016). Habitability: A Review. Astrobiology, 16(1), 89–117. 10.1089/ast.2015.1295

Colman, D. R., Kraus, E. A., Thieringer, P. H., Rempfert, K., Templeton, A. S., Spear, J. R., & Boyd, E. S. (2022). Deep-branching acetogens in serpentinized subsurface fluids of Oman. Proceedings of the National Academy of Sciences, 119(42), e2206845119. 10.1073/pnas.2206845119

Conrad, Ralf., & Seiler, Wolfgang. (1985). Characteristics of abiological carbon monoxide formation from soil organic matter, humic acids, and phenolic compounds. Environmental Science & Technology, 19(12), 1165–1169. 10.1021/es00142a004

Cook, M. C., Blank, J. G., Rietze, A., Suzuki, S., Nealson, K. H., & Morrill, P. L. (2021). A Geochemical Comparison of Three Terrestrial Sites of Serpentinization: The Tablelands, the Cedars, and Aqua de Ney. Journal of Geophysical Research: Biogeosciences, 126(11), e2021JG006316. 10.1029/2021JG006316

Cook, M. C., Blank, J. G., Suzuki, S., Nealson, K. H., & Morrill, P. L. (2021). Assessing Geochemical Bioenergetics and Microbial Metabolisms at Three Terrestrial Sites of Serpentinization: The Tablelands (NL, CAN), The Cedars (CA, USA), and Aqua de Ney (CA, USA). Journal of Geophysical Research: Biogeosciences, 126(6), e2019JG005542. 10.1029/2019JG005542

Costa, E., Pérez, J., & Kreft, J. U. (2006). Why is metabolic labour divided in nitrification? Trends in Microbiology, 14(5), 213–219. 10.1016/j.tim.2006.03.006

Crespo-Medina, M., Twing, K. I., Kubo, M. D. Y., Hoehler, T. M., Cardace, D., McCollom, T., & Schrenk, M. O. (2014). Insights into environmental controls on microbial communities in a continental serpentinite aquifer using a microcosm-based approach. Frontiers in Microbiology, 5. 10.3389/fmicb.2014.00604

Daims, H., Lebedeva, E. V., Pjevac, P., Han, P., Herbold, C., Albertsen, M., et al. (2015). Complete nitrification by Nitrospira bacteria. Nature, 528(7583), 504–509. 10.1038/nature16461

Daniels, L., Fuchs, G., Thauer, R. K., & Zeikus, J. G. (1977). Carbon Monoxide Oxidation by Methanogenic Bacteria. Journal of Bacteriology, 132(1), 118–126. 10.1128/jb.132.1.118-126.1977

Dick, J. M. (2019). CHNOSZ: Thermodynamic Calculations and Diagrams for Geochemistry. Frontiers in Earth Science, 0, 180. 10.3389/FEART.2019.00180

Ehlmann, B. L., Mustard, J. F., & Murchie, S. L. (2010). Geologic setting of serpentine deposits on Mars. Geophysical Research Letters, 37(6). 10.1029/2010GL042596

Eickenbusch, P., Takai, K., Sissman, O., Suzuki, S., Menzies, C., Sakai, S., et al. (2019). Origin of Short-Chain Organic Acids in Serpentinite Mud Volcanoes of the Mariana Convergent Margin. Frontiers in Microbiology, 10. 10.3389/fmicb.2019.01729

Gihring, T. M., Moser, D. P., Lin, L.-H., Davidson, M., Onstott, T. C., Morgan, L., et al. (2006). The Distribution of Microbial Taxa in the Subsurface Water of the Kalahari Shield, South Africa. Geomicrobiology Journal, 23(6), 415–430. 10.1080/01490450600875696

Glein, C. R., & Truong, N. (2025). Phosphates Reveal High pH Ocean Water on Enceladus. Icarus, 441, 116717. 10.1016/j.icarus.2025.116717

Glein, C. R., Baross, J. A., & Waite, J. H. (2015). The pH of Enceladus’ ocean. Geochimica et Cosmochimica Acta, 162, 202–219. 10.1016/j.gca.2015.04.017

Halevy, I., & Bachan, A. (2017). The geologic history of seawater pH. Science, 355(6329), 1069–1071. 10.1126/science.aal4151

Hand, K. P., Carlson, R. W., & Chyba, C. F. (2007). Energy, Chemical Disequilibrium, and Geological Constraints on Europa. Astrobiology, 7(6), 1006–1022. 10.1089/ast.2007.0156

Hashimoto, G. L., Abe, Y., & Sugita, S. (2007). The chemical composition of the early terrestrial atmosphere: Formation of a reducing atmosphere from CI-like material. Journal of Geophysical Research: Planets, 112(E5). 10.1029/2006JE002844

Helgeson, H. C., Kirkham, D. H., & Flowers, G. C. (1981). Theoretical prediction of the thermodynamic behavior of aqueous electrolytes by high pressures and temperatures; IV, Calculation of activity coefficients, osmotic coefficients, and apparent molal and standard and relative partial molal properties to 600 degrees C and 5kb. American Journal of Science, 281(10), 1249–1516. 10.2475/ajs.281.10.1249

Henstra, A. M., & Stams, A. J. M. (2011). Deep Conversion of Carbon Monoxide to Hydrogen and Formation of Acetate by the Anaerobic Thermophile Carboxydothermus hydrogenoformans. International Journal of Microbiology, 2011(1), 641582. 10.1155/2011/641582

Hinrichs, K.-U., Hayes, J. M., Sylva, S. P., Brewer, P. G., & DeLong, E. F. (1999). Methane-consuming archaebacteria in marine sediments. Nature, 398(6730), 802–805. 10.1038/19751

Hoefen, T. M., Clark, R. N., Bandfield, J. L., Smith, M. D., Pearl, J. C., & Christensen, P. R. (2003). Discovery of Olivine in the Nili Fossae Region of Mars. Science, 302(5645), 627–630. 10.1126/science.1089647

Hoehler, T. M. (2007). An Energy Balance Concept for Habitability. Astrobiology, 7(6), 824–838. 10.1089/ast.2006.0095

Hoehler, T. M., Amend, J. P., & Shock, E. L. (2007). A “Follow the Energy” Approach for Astrobiology. Astrobiology, 7(6), 819–823. 10.1089/ast.2007.0207

Holland, G., Lollar, B. S., Li, L., Lacrampe-Couloume, G., Slater, G. F., & Ballentine, C. J. (2013). Deep fracture fluids isolated in the crust since the Precambrian era. Nature, 497(7449), 357–360. 10.1038/nature12127

Holland, H. D. (1962). Model for the Evolution of the Earth’s Atmosphere. In A. E. J. Engel, H. L. James, & B. F. Leonard (Eds.), Petrologic Studies (p. 0). Geological Society of America. 10.1130/Petrologic.1962.447

Holland, H. D. (1984). The Chemical Evolution of the Atmosphere and Oceans. Princeton University Press.

Horita, J., & Berndt, M. E. (1999). Abiogenic Methane Formation and Isotopic Fractionation Under Hydrothermal Conditions. Science, 285(5430), 1055–1057. 10.1126/science.285.5430.1055

Howells, A. E. G., Quinn, L. M., Silva, M. G., Akiyama, K., Fifer, L. M., Boyer, G., et al. (2025). Energetic and genomic potential for hydrogenotrophic, formatotrophic, and acetoclastic methanogenesis in surface-expressed serpentinized fluids of the Samail Ophiolite. Frontiers in Microbiology, 15. 10.3389/fmicb.2024.1523912

Hu, R., Aronson, H. S., Weaver, M. E., Price, M. N., LaRowe, D. E., Liang, Y., et al. (2025, May 13). Organic carbon oxidation state shapes fermentative methanogenic microbiomes and controls greenhouse gas fluxes. bioRxiv. 10.1101/2025.05.12.653603

Hutchins, D. A., & Capone, D. G. (2022). The marine nitrogen cycle: new developments and global change. Nature Reviews Microbiology, 20(7), 401–414. 10.1038/s41579-022-00687-z

Jakosky, B. M., & Shock, E. L. (1998). The biological potential of Mars, the early Earth, and Europa. Journal of Geophysical Research: Planets, 103(E8), 19359–19364. 10.1029/98JE01892

Jones, R. M., Goordial, J. M., & Orcutt, B. N. (2018). Low Energy Subsurface Environments as Extraterrestrial Analogs. Frontiers in Microbiology, 9. 10.3389/fmicb.2018.01605

van Kessel, M. A. H. J., Speth, D. R., Albertsen, M., Nielsen, P. H., Op den Camp, H. J. M., Kartal, B., et al. (2015). Complete nitrification by a single microorganism. Nature, 528(7583), 555–559. 10.1038/nature16459

Kharecha, P., Kasting, J., & Siefert, J. (2005). A coupled atmosphere–ecosystem model of the early Archean Earth. Geobiology, 3(2), 53–76. 10.1111/j.1472-4669.2005.00049.x

Kieft, T. L., McCuddy, S. M., Onstott, T. C., Davidson, M., Lin, L.-H., Mislowack, B., et al. (2005). Geochemically Generated, Energy-Rich Substrates and Indigenous Microorganisms in Deep, Ancient Groundwater. Geomicrobiology Journal, 22(6), 325–335. 10.1080/01490450500184876

Knittel, K., & Boetius, A. (2009). Anaerobic Oxidation of Methane: Progress with an Unknown Process. Annual Review of Microbiology, 63(Volume 63, 2009), 311–334. 10.1146/annurev.micro.61.080706.093130

Koyama, S., Kamada, A., Furukawa, Y., Terada, N., Nakamura, Y., Yoshida, T., et al. (2024). Atmospheric formaldehyde production on early Mars leading to a potential formation of bio-important molecules. Scientific Reports, 14(1), 2397. 10.1038/s41598-024-52718-9

Kremp, F., Poehlein, A., Daniel, R., & Müller, V. (2018). Methanol metabolism in the acetogenic bacterium Acetobacterium woodii. Environmental Microbiology, 20(12), 4369–4384. 10.1111/1462-2920.14356

Lahijani, P., Zainal, Z. A., Mohammadi, M., & Mohamed, A. R. (2015). Conversion of the greenhouse gas CO2 to the fuel gas CO via the Boudouard reaction: A review. Renewable and Sustainable Energy Reviews, 41, 615–632. 10.1016/j.rser.2014.08.034

LaRowe, D., & Amend, J. (2014). Energetic constraints on life in marine deep sediments. In J. Kallmeyer & D. Wagner (Eds.), Microbial Life of the Deep Biosphere (pp. 279–302). DE GRUYTER. 10.1515/9783110300130.279

LaRowe, D., & Amend, J. (2019). Energy Limits for Life in the Subsurface. In B. N. Orcutt, I. Daniel, & R. Dasgupta (Eds.), Deep Carbon: Past to Present (pp. 585–619). Cambridge: Cambridge University Press. Retrieved from https://www.cambridge.org/core/books/deep-carbon/energy-limits-for-life-in-the-subsurface/A75173D093B9A43C6237C4F61821FDA8

LaRowe, D. E., & Amend, J. P. (2019). The Energetics of Fermentation in Natural Settings. Geomicrobiology Journal, 36(6), 492–505. 10.1080/01490451.2019.1573278

LaRowe, D. E., & Van Cappellen, P. (2011). Degradation of natural organic matter: A thermodynamic analysis. Geochimica et Cosmochimica Acta, 75(8), 2030–2042. 10.1016/j.gca.2011.01.020

LaRowe, D. E., Carlson, H. K., & Amend, J. P. (2021). The Energetic Potential for Undiscovered Manganese Metabolisms in Nature. Frontiers in Microbiology, 12. Retrieved from https://www.frontiersin.org/articles/10.3389/fmicb.2021.636145

Lau, M. C. Y., Kieft, T. L., Kuloyo, O., Linage-Alvarez, B., Van Heerden, E., Lindsay, M. R., et al. (2016). An oligotrophic deep-subsurface community dependent on syntrophy is dominated by sulfur-driven autotrophic denitrifiers. Proceedings of the National Academy of Sciences, 113(49). 10.1073/pnas.1612244113

Leong, J. A. M., & Shock, E. L. (2020). Thermodynamic constraints on the geochemistry of low-temperature, continental, serpentinization-generated fluids. American Journal of Science, 320(3), 185–235. 10.2475/03.2020.01

Lever, M. A. (2012). Acetogenesis in the Energy-Starved Deep Biosphere – A Paradox? Frontiers in Microbiology, 2. 10.3389/fmicb.2011.00284

Litty, D., Kremp, F., & Müller, V. (2022). One substrate, many fates: different ways of methanol utilization in the acetogen. Environmental Microbiology, 24(7), 3124–3133. 10.1111/1462-2920.16011

Lizarralde, D., Teske, A., Höfig, T. W., González-Fernández, A., & the IODP Expedition 385 Scientists. (2023). Carbon released by sill intrusion into young sediments measured through scientific drilling. Geology, 51(4), 329–333. 10.1130/G50665.1

Lu, G.-S., LaRowe, D. E., Fike, D. A., Druschel, G. K., Iii, W. P. G., Price, R. E., & Amend, J. P. (2020). Bioenergetic characterization of a shallow-sea hydrothermal vent system: Milos Island, Greece. PLOS ONE, 15(6), e0234175. 10.1371/journal.pone.0234175

Magnabosco, C., Ryan, K., Lau, M. C. Y., Kuloyo, O., Sherwood Lollar, B., Kieft, T. L., et al. (2016). A metagenomic window into carbon metabolism at 3 km depth in Precambrian continental crust. The ISME Journal, 10(3), 730–741. 10.1038/ismej.2015.150

Mara, P., Geller-McGrath, D., Edgcomb, V., Beaudoin, D., Morono, Y., & Teske, A. (2023). Metagenomic profiles of archaea and bacteria within thermal and geochemical gradients of the Guaymas Basin deep subsurface. Nature Communications, 14(1), 7768. 10.1038/s41467-023-43296-x

Marlow, J. J., LaRowe, D. E., Ehlmann, B. L., Amend, J. P., & Orphan, V. J. (2014). The Potential for Biologically Catalyzed Anaerobic Methane Oxidation on Ancient Mars. Astrobiology, 14(4), 292–307. 10.1089/ast.2013.1078

Martin, W., Baross, J., Kelley, D., & Russell, M. J. (2008). Hydrothermal vents and the origin of life. Nature Reviews Microbiology, 6(11), 805–814. 10.1038/nrmicro1991

McCollom, T. M. (1999). Methanogenesis as a potential source of chemical energy for primary biomass production by autotrophic organisms in hydrothermal systems on Europa. Journal of Geophysical Research: Planets, 104(E12), 30729–30742. 10.1029/1999JE001126

McCollom, T. M. (2000). Geochemical constraints on primary productivity in submarine hydrothermal vent plumes. Deep-Sea Research Part I: Oceanographic Research Papers, 47(1), 85–101. 10.1016/S0967-0637(99)00048-5

McCollom, T. M., & Seewald, J. S. (2007). Abiotic Synthesis of Organic Compounds in Deep-Sea Hydrothermal Environments. Chemical Reviews, 107(2), 382–401. 10.1021/cr0503660

McCollom, T. M., & Seewald, J. S. (2013). Serpentinites, Hydrogen, and Life. Elements, 9(2), 129–134. 10.2113/gselements.9.2.129

Merino, N., Aronson, H. S., Bojanova, D. P., Feyhl-Buska, J., Wong, M. L., Zhang, S., & Giovannelli, D. (2019). Living at the Extremes: Extremophiles and the Limits of Life in a Planetary Context. Frontiers in Microbiology, 10. 10.3389/fmicb.2019.00780

Michalski, J. R., Cuadros, J., Niles, P. B., Parnell, J., Deanne Rogers, A., & Wright, S. P. (2013). Groundwater activity on Mars and implications for a deep biosphere. Nature Geoscience, 6(2), 133–138. 10.1038/ngeo1706

Michalski, J. R., Onstott, T. C., Mojzsis, S. J., Mustard, J., Chan, Q. H. S., Niles, P. B., & Johnson, S. S. (2018). The Martian subsurface as a potential window into the origin of life. Nature Geoscience, 11(1), 21–26. 10.1038/s41561-017-0015-2

Momper, L., Casar, C. P., & Osburn, M. R. (2023). A metagenomic view of novel microbial and metabolic diversity found within the deep terrestrial biosphere at DeMMO: A microbial observatory in South Dakota, USA. Environmental Microbiology, 25(12), 3719–3737. 10.1111/1462-2920.16543

Morrill, P. L., Brazelton, W. J., Kohl, L., Rietze, A., Miles, S. M., Kavanagh, H., et al. (2014). Investigations of potential microbial methanogenic and carbon monoxide utilization pathways in ultra-basic reducing springs associated with present-day continental serpentinization: the Tablelands, NL, CAN. Frontiers in Microbiology, 5. 10.3389/fmicb.2014.00613

Moser, D. P., Gihring, T. M., Brockman, F. J., Fredrickson, J. K., Balkwill, D. L., Dollhopf, M. E., et al. (2005). Desulfotomaculum and Methanobacterium spp. Dominate a 4-to 5-Kilometer-Deep Fault. Applied and Environmental Microbiology, 71(12), 8773–8783. 10.1128/AEM.71.12.8773-8783.2005

Nitschke, W., & Russell, M. J. (2013). Beating the acetyl coenzyme A-pathway to the origin of life. Philosophical Transactions of the Royal Society B: Biological Sciences, 368(1622), 20120258. 10.1098/rstb.2012.0258

Nobu, M. K., Nakai, R., Tamazawa, S., Mori, H., Toyoda, A., Ijiri, A., et al. (2023). Unique H2-utilizing lithotrophy in serpentinite-hosted systems. The ISME Journal, 17(1), 95–104. 10.1038/s41396-022-01197-9

Novikov, A. A., Sokolova, T. G., Lebedinsky, A. V., Kolganova, T. V., & Bonch-Osmolovskaya, E. A. (2011). Carboxydothermus islandicus sp. nov., a thermophilic, hydrogenogenic, carboxydotrophic bacterium isolated from a hot spring. International Journal of Systematic and Evolutionary Microbiology, 61(10), 2532–2537. 10.1099/ijs.0.030288-0

Nyyssönen, M., Bomberg, M., Kapanen, A., Nousiainen, A., Pitkänen, P., & Itävaara, M. (2012). Methanogenic and Sulphate-Reducing Microbial Communities in Deep Groundwater of Crystalline Rock Fractures in Olkiluoto, Finland. Geomicrobiology Journal, 29(10), 863–878. 10.1080/01490451.2011.635759

O’Brien, J. M., Wolkin, R. H., Moench, T. T., Morgan, J. B., & Zeikus, J. G. (1984). Association of hydrogen metabolism with unitrophic or mixotrophic growth of Methanosarcina barkeri on carbon monoxide. Journal of Bacteriology, 158(1), 373–375. 10.1128/jb.158.1.373-375.1984

Ody, A., Poulet, F., Langevin, Y., Bibring, J.-P., Bellucci, G., Altieri, F., et al. (2012). Global maps of anhydrous minerals at the surface of Mars from OMEGA/MEx. Journal of Geophysical Research: Planets, 117(E11). 10.1029/2012JE004117

Oelgeschläger, E., & Rother, M. (2008). Carbon monoxide-dependent energy metabolism in anaerobic bacteria and archaea. Archives of Microbiology, 190(3), 257–269. 10.1007/s00203-008-0382-6

Onstott, T. C., McGown, D. J., Bakermans, C., Ruskeeniemi, T., Ahonen, L., Telling, J., et al. (2009). Microbial Communities in Subpermafrost Saline Fracture Water at the Lupin Au Mine, Nunavut, Canada. Microbial Ecology, 58(4), 786–807. 10.1007/s00248-009-9553-5

Orphan, V. J., House, C. H., Hinrichs, K.-U., McKeegan, K. D., & DeLong, E. F. (2001). Methane-Consuming Archaea Revealed by Directly Coupled Isotopic and Phylogenetic Analysis. Science, 293(5529), 484–487. 10.1126/science.1061338

Osburn, M. R., LaRowe, D. E., Momper, L. M., & Amend, J. P. (2014). Chemolithotrophy in the continental deep subsurface: Sanford Underground Research Facility (SURF), USA. Frontiers in Microbiology, 5. 10.3389/fmicb.2014.00610

Osburn, M. R., Kruger, B., Masterson, A. L., Casar, C. P., & Amend, J. P. (2019). Establishment of the Deep Mine Microbial Observatory (DeMMO), South Dakota, USA, a Geochemically Stable Portal Into the Deep Subsurface. Frontiers in Earth Science, 7. 10.3389/feart.2019.00196

Ozaki, K., Tajika, E., Hong, P. K., Nakagawa, Y., & Reinhard, C. T. (2018). Effects of primitive photosynthesis on Earth’s early climate system. Nature Geoscience, 11(1), 55–59. 10.1038/s41561-017-0031-2

Parshina, S. N., Sipma, J., Nakashimada, Y., Henstra, A. M., Smidt, H., Lysenko, A. M., et al. (2005). Desulfotomaculum carboxydivorans sp. nov., a novel sulfate-reducing bacterium capable of growth at 100 % CO. International Journal of Systematic and Evolutionary Microbiology, 55(5), 2159–2165. 10.1099/ijs.0.63780-0

Parshina, S. N., Sipma, J., Henstra, A. M., & Stams, A. J. M. (2010). Carbon Monoxide as an Electron Donor for the Biological Reduction of Sulphate. International Journal of Microbiology, 2010(1), 319527. 10.1155/2010/319527

Poole, J. H. J. (1951). The evolution of the earth’s atmosphere. Sci. Proc. R. Dublin Soc., 25, 201–224.

Qi, Q.-C., Meng, N., Li, S., Wang, J., Ran, X., & Zhuang, G.-C. (2025). Occurrence and Cycling of Carbon Monoxide in Marine Coastal Sediments. Journal of Geophysical Research: Oceans, 130(4), e2025JC022439. 10.1029/2025JC022439

Ragsdale, S. W., & Pierce, E. (2008). Acetogenesis and the Wood–Ljungdahl pathway of CO2 fixation. Biochimica et Biophysica Acta (BBA) - Proteins and Proteomics, 1784(12), 1873–1898. 10.1016/j.bbapap.2008.08.012

Robare, J. (2025). Thermodynamic Modeling of Chemical Energy Supplies and Demands of Geobiochemical Systems (Ph.D.). Arizona State University, United States -- Arizona. Retrieved from https://www.proquest.com/docview/3230363409/abstract/E5CFE17B9C2E4059PQ/1

Rother, M., & Metcalf, W. W. (2004). Anaerobic growth of *Methanosarcina acetivorans* C2A on carbon monoxide: An unusual way of life for a methanogenic archaeon. Proceedings of the National Academy of Sciences, 101(48), 16929–16934. 10.1073/pnas.0407486101

Rucker, H. R., Ely, T. D., LaRowe, D. E., Giovannelli, D., & Price, R. E. (2023). Quantifying the Bioavailable Energy in an Ancient Hydrothermal Vent on Mars and a Modern Earth-Based Analog. Astrobiology, 23(4), 431–445. 10.1089/ast.2022.0064

Russell, M. J., Hall, A. J., & Martin, W. (2010). Serpentinization as a source of energy at the origin of life. Geobiology, 8(5), 355–371. 10.1111/j.1472-4669.2010.00249.x

Russell, Michael J., & Martin, W. (2004). The rocky roots of the acetyl-CoA pathway. Trends in Biochemical Sciences, 29(7), 358–363. 10.1016/j.tibs.2004.05.007

Russell, Michael J., & Nitschke, W. (2017). Methane: Fuel or Exhaust at the Emergence of Life? Astrobiology, 17(10), 1053–1066. 10.1089/ast.2016.1599

Sabuda, M. C., Putman, L. I., Hoehler, T. M., Kubo, M. D., Brazelton, W. J., Cardace, D., & Schrenk, M. O. (2021). Biogeochemical Gradients in a Serpentinization-Influenced Aquifer: Implications for Gas Exchange Between the Subsurface and Atmosphere. Journal of Geophysical Research: Biogeosciences, 126(8), e2020JG006209. 10.1029/2020JG006209

Schaefer, L., & Fegley, B. (2007). Outgassing of ordinary chondritic material and some of its implications for the chemistry of asteroids, planets, and satellites. Icarus, 186(2), 462–483. 10.1016/j.icarus.2006.09.002

Schaefer, L., & Fegley, B. (2010). Chemistry of atmospheres formed during accretion of the Earth and other terrestrial planets. Icarus, 208(1), 438–448. 10.1016/j.icarus.2010.01.026

Seewald, J. S., Zolotov, M. Yu., & McCollom, T. (2005). Experimental investigation of single carbon compounds under hydrothermal conditions. Geochimica et Cosmochimica Acta, 70(2), 446–460. 10.1016/j.gca.2005.09.002

Sherwood Lollar, B., Frape, S. K., Weise, S. M., Fritz, P., Macko, S. A., & Welhan, J. A. (1993). Abiogenic methanogenesis in crystalline rocks. Geochimica et Cosmochimica Acta, 57(23), 5087–5097. 10.1016/0016-7037(93)90610-9

Sherwood Lollar, B., Lacrampe-Couloume, G., Slater, G. F., Ward, J., Moser, D. P., Gihring, T. M., et al. (2006). Unravelling abiogenic and biogenic sources of methane in the Earth’s deep subsurface. Chemical Geology, 226(3), 328–339. 10.1016/j.chemgeo.2005.09.027

Shock, E. L. (1997). High-temperature life without photosynthesis as a model for Mars. Journal of Geophysical Research: Planets, 102(E10), 23687–23694. 10.1029/97JE01087

Shock, E. L., & Helgeson, H. C. (1988). Calculation of the thermodynamic and transport properties of aqueous species at high pressures and temperatures: Correlation algorithms for ionic species and equation of state predictions to 5 kb and 1000°C. Geochimica et Cosmochimica Acta, 52(8), 2009–2036. 10.1016/0016-7037(88)90181-0

Shock, E. L., & Holland, M. E. (2007). Quantitative Habitability. Astrobiology, 7(6), 839–851. 10.1089/ast.2007.0137

Shock, E. L., Holland, M., Meyer-Dombard, D., Amend, J. P., Osburn, G. R., & Fischer, T. P. (2010). Quantifying inorganic sources of geochemical energy in hydrothermal ecosystems, Yellowstone National Park, USA. Geochimica et Cosmochimica Acta, 74(14), 4005–4043. 10.1016/j.gca.2009.08.036

Slepova, T. V., Sokolova, T. G., Lysenko, A. M., Tourova, T. P., Kolganova, T. V., Kamzolkina, O. V., et al. (2006). Carboxydocella sporoproducens sp. nov., a novel anaerobic CO-utilizing/H2-producing thermophilic bacterium from a Kamchatka hot spring. International Journal of Systematic and Evolutionary Microbiology, 56(4), 797–800. 10.1099/ijs.0.63961-0

Slepova, T. V., Sokolova, T. G., Kolganova, T. V., Tourova, T. P., & Bonch-Osmolovskaya, E. A. (2009). Carboxydothermus siderophilus sp. nov., a thermophilic, hydrogenogenic, carboxydotrophic, dissimilatory Fe(III)-reducing bacterium from a Kamchatka hot spring. International Journal of Systematic and Evolutionary Microbiology, 59(2), 213–217. 10.1099/ijs.0.000620-0

Sokolova, T G, González, J. M., Kostrikina, N. A., Chernyh, N. A., Tourova, T. P., Kato, C., et al. (2001). Carboxydobrachium pacificum gen. nov., sp. nov., a new anaerobic, thermophilic, CO-utilizing marine bacterium from Okinawa Trough. International Journal of Systematic and Evolutionary Microbiology, 51(1), 141–149. 10.1099/00207713-51-1-141

Sokolova, Tatyana G., González, J. M., Kostrikina, N. A., Chernyh, N. A., Slepova, T. V., Bonch-Osmolovskaya, E. A., & Robb, F. T. (2004). Thermosinus carboxydivorans gen. nov., sp. nov., a new anaerobic, thermophilic, carbon-monoxide-oxidizing, hydrogenogenic bacterium from a hot pool of Yellowstone National Park. International Journal of Systematic and Evolutionary Microbiology, 54(6), 2353–2359. 10.1099/ijs.0.63186-0

Steiger, M. G., Mattanovich, D., & Sauer, M. (2017). Microbial organic acid production as carbon dioxide sink. FEMS Microbiology Letters, 364(21). 10.1093/femsle/fnx212

Suzuki, S., Nealson, K. H., & Ishii, S. (2018). Genomic and in-situ Transcriptomic Characterization of the Candidate Phylum NPL-UPL2 From Highly Alkaline Highly Reducing Serpentinized Groundwater. Frontiers in Microbiology, 9. 10.3389/fmicb.2018.03141

Suzuki, S., Ishii, S., Chadwick, G. L., Tanaka, Y., Kouzuma, A., Watanabe, K., et al. (2024). A non-methanogenic archaeon within the order Methanocellales. Nature Communications, 15(1), 4858. 10.1038/s41467-024-48185-5

Svetlichny, V. A., Sokolova, T. G., Gerhardt, M., Ringpfeil, M., Kostrikina, N. A., & Zavarzin, G. A. (1991). *Carboxydothermus hydrogenoformans* gen. nov., sp. nov., a CO-utilizing Thermophilic Anaerobic Bacterium from Hydrothermal Environments of Kunashir Island. Systematic and Applied Microbiology, 14(3), 254–260. 10.1016/S0723-2020(11)80377-2

Tanger, J. C., & Helgeson, H. C. (1988). Calculation of the thermodynamic and transport properties of aqueous species at high pressure and temperatures; revised equations of state for the standard partial molal properties of ions and electrolytes. American Journal of Science, 288(1), 19–98.

Twing, K. I., Brazelton, W. J., Kubo, M. D. Y., Hyer, A. J., Cardace, D., Hoehler, T. M., et al. (2017). Serpentinization-Influenced Groundwater Harbors Extremely Low Diversity Microbial Communities Adapted to High pH. Frontiers in Microbiology, 8. Retrieved from https://www.frontiersin.org/articles/10.3389/fmicb.2017.00308

Urey, H. C. (1952). On the Early Chemical History of the Earth and the Origin of Life. Proceedings of the National Academy of Sciences, 38(4), 351–363. 10.1073/pnas.38.4.351

Van de Graaf, A. A., Mulder, A., De Bruijn, P., Jetten, M. S. M., Robertson, L. A., & Kuenen, J. G. (1995). Anaerobic oxidation of ammoinium is a biologically mediated process. Applied and Environmental Microbiology, 61(4), 1246–1251. PMC167380

Vance, S., Harnmeijer, J., Kimura, J., Hussmann, H., deMartin, B., & Brown, J. M. (2007). Hydrothermal Systems in Small Ocean Planets. Astrobiology, 7(6), 987–1005. 10.1089/ast.2007.0075

Vance, S. D., & Melwani Daswani, M. (2020). Serpentinite and the search for life beyond Earth. *Philosophical Transactions of the Royal Society A: Mathematical*, Physical and Engineering Sciences, 378(2165), 20180421. 10.1098/rsta.2018.0421

Vance, S. D., Hand, K. P., & Pappalardo, R. T. (2016). Geophysical controls of chemical disequilibria in Europa. Geophysical Research Letters, 43(10), 4871–4879. 10.1002/2016GL068547

Vick, T., Dodsworth, J., Costa, K., Shock, E. L., & Hedlund, B. P. (2010). Microbiology and geochemistry of Little Hot Creek, a hot spring environment in the Long Valley Caldera. Geobiology, 8, 140–154.

Von Damm, K. L., Edmond, J. M., Measures, C. I., & Grant, B. (1985). Chemistry of submarine hydrothermal solutions at Guaymas Basin, Gulf of California. Geochimica et Cosmochimica Acta, 49(11), 2221–2237. 10.1016/0016-7037(85)90223-6

Waite, J. H., Glein, C. R., Perryman, R. S., Teolis, B. D., Magee, B. A., Miller, G., et al. (2017). Cassini finds molecular hydrogen in the Enceladus plume: Evidence for hydrothermal processes. Science, 356(6334), 155–159. 10.1126/science.aai8703

Weiss, M. C., Sousa, F. L., Mrnjavac, N., Neukirchen, S., Roettger, M., Nelson-Sathi, S., & Martin, W. F. (2016). The physiology and habitat of the last universal common ancestor. Nature Microbiology, 1(9), 16116. 10.1038/nmicrobiol.2016.116

Wogan, N. F., Catling, D. C., Zahnle, K. J., & Lupu, R. (2023). Origin-of-life Molecules in the Atmosphere after Big Impacts on the Early Earth. The Planetary Science Journal, 4(9), 169. 10.3847/PSJ/aced83

Yanez, M. D., LaRowe, D. E., Cable, M. L., & Amend, J. P. (2024). Energy yields for acetylenotrophy on Enceladus and Titan. Icarus, 411, 115969. 10.1016/j.icarus.2024.115969

Zahnle, K., Schaefer, L., & Fegley, B. (2010). Earth’s Earliest Atmospheres. Cold Spring Harbor Perspectives in Biology, 2(10), a004895–a004895. 10.1101/cshperspect.a004895

Zolotov, M. Y., & Shock, E. L. (2003). Energy for biologic sulfate reduction in a hydrothermally formed ocean on Europa. Journal of Geophysical Research: Planets, 108(E4). 10.1029/2002JE001966

Zolotov, M. Y., & Shock, E. L. (2004). A model for low-temperature biogeochemistry of sulfur, carbon, and iron on Europa. Journal of Geophysical Research: Planets, 109(E6). 10.1029/2003JE002194

## References

Accornero, M., Marini, L., & Lelli, M. (2010). Prediction of the thermodynamic properties of metal–chromate aqueous complexes to high temperatures and pressures and implications for the speciation of hexavalent chromium in some natural waters. Applied Geochemistry, 25(2), 242–260. 10.1016/j.apgeochem.2009.11.010

Akilan, C., Rohman, N., Hefter, G., & Buchner, R. (2006). Temperature Effects on Ion Association and Hydration in MgSO4 by Dielectric Spectroscopy. ChemPhysChem, 7(11), 2319–2330. 10.1002/cphc.200600342

Akinfiev, N. N., & Tagirov, B. R. (2014). Zn in hydrothermal systems: Thermodynamic description of hydroxide, chloride, and hydrosulfide complexes. Geochemistry International, 52(3), 197–214. 10.1134/S0016702914030021

Akinfiev, N. N., & Zotov, A. V. (2010). Thermodynamic description of aqueous species in the system Cu-Ag-Au-S-O-H at temperatures of 0–600°C and pressures of 1–3000 bar. Geochemistry International, 48(7), 714–720. 10.1134/S0016702910070074

Akinfiev, N. N., & Zotov, A. V. (2023). Copper in Hydrothermal Systems: a Thermodynamic Description of Hydroxocomplexes. Geology of Ore Deposits, 65(1), 1–10. 10.1134/S1075701523010026

Akinfiev, N. N., Voronin, M. V., Zotov, A. V., & Prokof’ev, V. Yu. (2006). Experimental investigation of the stability of a chloroborate complex and thermodynamic description of aqueous species in the B-Na-Cl-O-H system up to 350°C. Geochemistry International, 44(9), 867–878. 10.1134/S0016702906090035

Akinfiev, Nikolay N., & Diamond, L. W. (2003). Thermodynamic description of aqueous nonelectrolytes at infinite dilution over a wide range of state parameters. Geochimica et Cosmochimica Acta, 67(4), 613–629. 10.1016/S0016-7037(02)01141-9

Akinfiev, Nikolay N., & Plyasunov, A. V. (2014). Application of the Akinfiev–Diamond equation of state to neutral hydroxides of metalloids (B(OH)3, Si(OH)4, As(OH)3) at infinite dilution in water over a wide range of the state parameters, including steam conditions. Geochimica et Cosmochimica Acta, 126, 338–351. 10.1016/j.gca.2013.11.013

Akinfiev, Nikolay N., Korzhinskaya, V. S., Kotova, N. P., Redkin, A. F., & Zotov, A. V. (2020). Niobium and tantalum in hydrothermal fluids: Thermodynamic description of hydroxide and hydroxofluoride complexes. Geochimica et Cosmochimica Acta, 280, 102–115. 10.1016/j.gca.2020.04.009

Akinfiev, N.N., & Zotov, A. V. (2001). Thermodynamic Description of Chloride, Hydrosulfide, and Hydroxo Complexes of Ag(I), Cu(I), and Au(I) at Temperatures of 25–500°C and Pressures of 1–2000 bar. Geochemistry International, 39(10), 990–1006. 10.1134/S0016702910070074

Amend, J P, & Shock, E. L. (2001). Energetics of overall metabolic reaction of thermophilic and hyperthermophylic Archaea and Bacteria. FEMS Microb. Revs., 25, 175–243.

Amend, Jan P., & Helgeson, H. C. (1997). Calculation of the standard molal thermodynamic properties of aqueous biomolecules at elevated temperatures and pressures Part1L-α-Amino acids. Journal of the Chemical Society, Faraday Transactions, 93(10), 1927–1941. 10.1039/A608126F

Amend, Jan P., & Plyasunov, A. V. (2001). Carbohydrates in thermophile metabolism: calculation of the standard molal thermodynamic properties of aqueous pentoses and hexoses at elevated temperatures and pressures. Geochimica et Cosmochimica Acta, 65(21), 3901–3917. 10.1016/S0016-7037(01)00707-4

Azadi, M. R., Karrech, A., Attar, M., & Elchalakani, M. (2019). Data analysis and estimation of thermodynamic properties of aqueous monovalent metal-glycinate complexes. Fluid Phase Equilibria, 480, 25–40. 10.1016/j.fluid.2018.10.002

Bandura, A. V., & Lvov, S. N. (2005). The Ionization Constant of Water over Wide Ranges of Temperature and Density. Journal of Physical and Chemical Reference Data, 35(1), 15–30. 10.1063/1.1928231

Barin, I., & Knacke, O. (1973). Thermochemical properties of inorganic substances. Berlin: Springer-Verlag.

Bazarkina, E. F., Zotov, A. V., & Akinfiev, N. N. (2010). Pressure-dependent stability of cadmium chloride complexes: Potentiometric measurements at 1–1000 bar and 25°C. Geology of Ore Deposits, 52(2), 167–178. 10.1134/S1075701510020054

Berman, R. G. (1988). Internally-Consistent Thermodynamic Data for Minerals in the System Na2O-K2O-CaO-MgO-FeO-Fe2O3-Al2O3-SiO2-TiO2-H2O-CO2. Journal of Petrology, 29(2), 445–522. 10.1093/petrology/29.2.445

Berman, R. G. (1990). Mixing properties of Ca-Mg-Fe-Mn garnets. American Mineralogist, 75(3–4), 328–344.

Bird, L. J., Bonnefoy, V., & Newman, D. K. (2011). Bioenergetic challenges of microbial iron metabolisms. Trends in Microbiology, 19(7), 330–340. 10.1016/j.tim.2011.05.001

Bowers, T. S., & Helgeson, H. C. (1983). Calculation of the thermodynamic and geochemical consequences of nonideal mixing in the system H2O-CO2-NaCl on phase relations in geologic systems: Equation of state for H2O-CO2-NaCl fluids at high pressures and temperatures. Geochimica et Cosmochimica Acta, 47(7), 1247–1275. 10.1016/0016-7037(83)90066-2

Boyer, G., & Shock, E. (2023, August 1). The Complicator (prototype). Zenodo. 10.5281/zenodo.8206644

Bradley, J. A., Amend, J. P., & LaRowe, D. E. (2019). Survival of the fewest: Microbial dormancy and maintenance in marine sediments through deep time. Geobiology, 17(1), 43–59. 10.1111/gbi.12313

Canovas, P. A., & Shock, E. L. (2016). Geobiochemistry of metabolism: Standard state thermodynamic properties of the citric acid cycle. Geochimica et Cosmochimica Acta, 195, 293–322. 10.1016/j.gca.2016.08.028

Dale, J. D., Shock, E. L., Macleod, G., Aplin, A. C., & Larter, S. R. (1997). Standard partial molal properties of aqueous alkylphenols at high pressures and temperatures. Geochimica et Cosmochimica Acta, 61(19), 4017–4024. 10.1016/S0016-7037(97)00212-3

Denbigh, K. G. (1981). The Principles of Chemical Equilibrium: With Applications in Chemistry and Chemical Engineering (4th ed.). Cambridge: Cambridge University Press. 10.1017/CBO9781139167604

Diakonov, I., Pokrovski, G., Schott, J., Castet, S., & Gout, R. (1996). An experimental and computational study of sodium-aluminum complexing in crustal fluids. Geochimica et Cosmochimica Acta, 60(2), 197–211. 10.1016/0016-7037(95)00403-3

Diakonov, I. I., Ragnarsdottir, K. V., & Tagirov, B. R. (1998). Standard thermodynamic properties and heat capacity equations of rare earth hydroxides:: II. Ce(III)-, Pr-, Sm-, Eu(III)-, Gd-, Tb-, Dy-, Ho-, Er-, Tm-, Yb-, and Y-hydroxides. Comparison of thermochemical and solubility data. Chemical Geology, 151(1), 327–347. 10.1016/S0009-2541(98)00088-6

Dick, J. M., LaRowe, D. E., & Helgeson, H. C. (2006). Temperature, pressure, and electrochemical constraints on protein speciation: Group additivity calculation of the standard molal thermodynamic properties of ionized unfolded proteins. Biogeosciences, 3(3), 311–336. 10.5194/bg-3-311-2006

Dick, Jeffrey M. (2019). CHNOSZ: Thermodynamic Calculations and Diagrams for Geochemistry. Frontiers in Earth Science, 0, 180. 10.3389/FEART.2019.00180

Dick, Jeffrey M., Evans, K. A., Holman, A. I., Jaraula, C. M. B., & Grice, K. (2013). Estimation and application of the thermodynamic properties of aqueous phenanthrene and isomers of methylphenanthrene at high temperature. Geochimica et Cosmochimica Acta, 122, 247–266. 10.1016/j.gca.2013.08.020

Dick, Jeffrey Michael. (2007). Calculation of the relative stabilities of proteins as a function of temperature, pressure, and chemical potentials in subcellular and geochemical environments (Ph.D.). University of California, Berkeley, United States -- California. Retrieved from https://www.proquest.com/docview/304901180/abstract/E8B2B32328F64C93PQ/1

Evans, B. W. (1990). Phase relations of epidote-blueschists. Lithos, 25(1), 3–23. 10.1016/0024-4937(90)90003-J

Facq, S., Daniel, I., Montagnac, G., Cardon, H., & Sverjensky, D. A. (2014). In situ Raman study and thermodynamic model of aqueous carbonate speciation in equilibrium with aragonite under subduction zone conditions. Geochimica et Cosmochimica Acta, 132, 375–390. 10.1016/j.gca.2014.01.030

GEOPIG. (2019). GEOPIG slop files [Data set]. Zenodo. 10.5281/zenodo.2630820

Goldberg, A. B., Maroulis, P. J., Wilner, L. A., & Bandy, A. R. (1981). Study of H2S emissions from a salt water marsh. Atmospheric Environment (1967), 15(1), 11–18. 10.1016/0004-6981(81)90119-0

Goldberg, R. N., Kishore, N., & Lennen, R. M. (2002). Thermodynamic Quantities for the Ionization Reactions of Buffers. Journal of Physical and Chemical Reference Data, 31(2), 231–370. 10.1063/1.1416902

Gottschalk, M. (2004). Thermodynamic Properties of Zoisite, Clinozoisite and Epidote. Reviews in Mineralogy and Geochemistry, 56(1), 83–124. 10.2138/gsrmg.56.1.83

Grenthe, I., Gaona, X., Plyasunov, A. V., Rao, L., Runde, W. H., Grambow, B., et al. (2021). Second update on the Chemical Thermodynamics of Uranium, Neptunium, Plutonium, Americium And Technetium, Volume 14. Chemical Thermodynamics, 14. 10.1787/bf86a907-en

Grevel, K.-D., & Majzlan, J. (2009). Internally consistent thermodynamic data for magnesium sulfate hydrates. Geochimica et Cosmochimica Acta, 73(22), 6805–6815. 10.1016/j.gca.2009.08.005

Gysi, A. P., Harlov, D., Filho, D. C., & Williams-Jones, A. E. (2016). Experimental determination of the high temperature heat capacity of a natural xenotime-(Y) solid solution and synthetic DyPO4 and ErPO4 endmembers. Thermochimica Acta, 627–629, 61–67. 10.1016/j.tca.2016.01.016

Haar, L., Gallagher, J. S., & Kell, G. S. (1984). NBS/NRC steam tables: thermodynamic and transport properties and computer programs for vapor and liquid states of water in SI units. Toronto: Hemisphere Pub. Corp.

Haas, J. R., & Shock, E. L. (1999). Halocarbons in the environment: estimates of thermodynamic properties for aqueous chloroethylene species and their stabilities in natural settings. Geochimica et Cosmochimica Acta, 63(19), 3429–3441. 10.1016/S0016-7037(99)00276-8

Haas, J. R., Shock, E. L., & Sassani, D. C. (1995). Rare earth elements in hydrothermal systems: Estimates of standard partial molal thermodynamic properties of aqueous complexes of the rare earth elements at high pressures and temperatures. Geochimica et Cosmochimica Acta, 59(21), 4329–4350. 10.1016/0016-7037(95)00314-P

Hakin, A. W., Duke, M. M., Marty, J. L., & Preuss, K. E. (1994). Some thermodynamic properties of aqueous amino acid systems at 288.15, 298.15, 313.15 and 328.15 K: group additivity analyses of standard-state volumes and heat capacities. Journal of the Chemical Society, Faraday Transactions, 90(14), 2027–2035. 10.1039/FT9949002027

Hawrylak, B., Palepu, R., & Tremaine, P. R. (2006). Thermodynamics of aqueous methyldiethanolamine (MDEA) and methyldiethanolammonium chloride (MDEAH+Cl−) over a wide range of temperature and pressure: Apparent molar volumes, heat capacities, and isothermal compressibilities. The Journal of Chemical Thermodynamics, 38(8), 988–1007. 10.1016/j.jct.2005.10.013

Helgeson, H. C. (1985). Errata II; Thermodynamics of minerals, reactions, and aqueous solutions at high pressures and temperatures. American Journal of Science, 285(9). 10.2475/ajs.285.9.845

Helgeson, H. C., Delany, J. M., Nesbitt, H. W., & Bird, D. K. (1978). Summary and critique of the thermodynamic properties of rock-forming minerals. American Journal of Science, 278-A.

Helgeson, H. C., Owens, C. E., Knox, A. M., & Richard, L. (1998). Calculation of the Standard Molal Thermodynamic Properties of Crystalline, Liquid, and Gas Organic Molecules at High Temperatures and Pressures. Geochimica et Cosmochimica Acta, 62(6), 985–1081. 10.1016/S0016-7037(97)00219-6

Helgeson, H. C., Richard, L., McKenzie, W. F., Norton, D. L., & Schmitt, A. (2009). A chemical and thermodynamic model of oil generation in hydrocarbon source rocks. Geochimica et Cosmochimica Acta, 73(3), 594–695. 10.1016/j.gca.2008.03.004

Hemingway, B. S. (1982). Thermodynamic properties of selected uranium compounds and aqueous species at 298.15 K and 1 bar and at higher temperatures; preliminary models for the origin of coginite deposits (No. 82–619). Open-File Report. U.S. Geological Survey,. 10.3133/ofr82619

Hemingway, Bruce S., Robie, R. A., & Apps, J. A. (1991). Revised values for the thermodynamic properties of boehmite, AIO(OH), and related species and phases in the system AI-H-O. American Mineralogist, 76, 445–457.

Ho, P. C., & Palmer, D. A. (1997). Ion association of dilute aqueous potassium chloride and potassium hydroxide solutions to 600°C and 300 MPa determined by electrical conductance measurements. Geochimica et Cosmochimica Acta, 61(15), 3027–3040. 10.1016/S0016-7037(97)00146-4

Hu, B., Etschmann, B., Testemale, D., Liu, W., Guan, Q., Müller, H., & Brugger, J. (2025). Tantalum in hydrothermal fluids. Geochimica et Cosmochimica Acta, 406, 243–261. 10.1016/j.gca.2024.10.019

Huang, F., & Sverjensky, D. A. (2019). Extended Deep Earth Water Model for predicting major element mantle metasomatism. Geochimica et Cosmochimica Acta, 254, 192–230. 10.1016/j.gca.2019.03.027

Huston, D. L., & Bastrakov, E. (2024). Germanium, indium, gallium and cadmium in zinc ores: a mineral system approach. Australian Journal of Earth Sciences, 71(8), 1125–1155. 10.1080/08120099.2024.2423772

Jackson, K. J., & Helgeson, H. C. (1985). Chemical and thermodynamic constraints on the hydrothermal transport and deposition of tin; II, Interpretation of phase relations in the Southeast Asian tin belt. Economic Geology, 80(5), 1365–1378. 10.2113/gsecongeo.80.5.1365

Jacob, K. T., Shekhar, C., & Waseda, Y. (2009). An update on the thermodynamics of Ta2O5. The Journal of Chemical Thermodynamics, 41(6), 748–753. 10.1016/j.jct.2008.12.006

Johnson, J. W., Oelkers, E. H., & Helgeson, H. C. (1992a). SUPCRT92: A software package for calculating the standard molal thermodynamic properties of minerals, gases, aqueous species, and reactions from 1 to 5000 bar and 0 to 1000°C. Computers & Geosciences, 18(7), 899–947. 10.1016/0098-3004(92)90029-Q

Johnson, J. W., Oelkers, E. H., & Helgeson, H. C. (1992b). SUPCRT92: A software package for calculating the standard molal thermodynamic properties of minerals, gases, aqueous species, and reactions from 1 to 5000 bar and 0 to 1000°C. Computers & Geosciences, 18(7), 899–947. 10.1016/0098-3004(92)90029-Q

Jørgensen, B. B., Findlay, A. J., & Pellerin, A. (2019). The Biogeochemical Sulfur Cycle of Marine Sediments. Frontiers in Microbiology, 10. 10.3389/fmicb.2019.00849

Kelley, K. K. (1960). Contributions to the data on theoretical metallurgy. XIII, High-temperature heat-content, heat-capacity, and entropy data for the elements and inorganic compounds. Washington D.C.: U.S. Dept. of the Interior, Bureau of Mines : G.P.O. Retrieved from http://digital.library.unt.edu/ark:/67531/metadc12739

Kitadai, N. (2014). Thermodynamic Prediction of Glycine Polymerization as a Function of Temperature and pH Consistent with Experimentally Obtained Results. Journal of Molecular Evolution, 78(3), 171–187. 10.1007/s00239-014-9616-1

Kitadai, N. (2015). Energetics of Amino Acid Synthesis in Alkaline Hydrothermal Environments. Origins of Life and Evolution of Biospheres, 45(4), 377–409. 10.1007/s11084-015-9428-3

Knight, A. W., Bryan, C. R., & Jové-Colón, C. F. (2024). Development of a consistent geochemical model of the Mg(OH)2–MgCl2–H2O system from 25°C to 120°C. Applied Geochemistry, 169, 106032. 10.1016/j.apgeochem.2024.106032

Konings, R.J.M., & Kovács, A. (2003). Thermodynamic properties of the lanthanide(III) halides. In Handbook on the Physics and Chemistry of Rare Earths (Vol. 33, pp. 147–247). Elsevier. 10.1016/S0168-1273(02)33003-4

Konings, Rudy J. M., Beneš, O., Kovács, A., Manara, D., Sedmidubský, D., Gorokhov, L., et al. (2014). The Thermodynamic Properties of the f-Elements and their Compounds. Part 2. The Lanthanide and Actinide Oxides. Journal of Physical and Chemical Reference Data, 43(1), 013101. 10.1063/1.4825256

Langmuir, D., Mahoney, J., & Rowson, J. (2006). Solubility products of amorphous ferric arsenate and crystalline scorodite (FeAsO4 · 2H2O) and their application to arsenic behavior in buried mine tailings. Geochimica et Cosmochimica Acta, 70(12), 2942–2956. 10.1016/j.gca.2006.03.006

LaRowe, D. E., & Amend, J. P. (2016). The energetics of anabolism in natural settings. The ISME Journal, 10(6), 1285–1295. 10.1038/ismej.2015.227

LaRowe, D. E., & Dick, J. M. (2012). Calculation of the standard molal thermodynamic properties of crystalline peptides. Geochimica et Cosmochimica Acta, 80, 70–91. 10.1016/j.gca.2011.11.041

LaRowe, D. E., & Helgeson, H. C. (2006a). Biomolecules in hydrothermal systems: Calculation of the standard molal thermodynamic properties of nucleic-acid bases, nucleosides, and nucleotides at elevated temperatures and pressures. Geochimica et Cosmochimica Acta, 70(18), 4680–4724. 10.1016/j.gca.2006.04.010

LaRowe, D. E., & Helgeson, H. C. (2006b). The energetics of metabolism in hydrothermal systems: Calculation of the standard molal thermodynamic properties of magnesium-complexed adenosine nucleotides and NAD and NADP at elevated temperatures and pressures. Thermochimica Acta, 448(2), 82–106. 10.1016/j.tca.2006.06.008

Laskar, C., Bazarkina, E. F., & Pokrovski, G. S. (2025). The role of sulfur in palladium transport and fractionation from platinum by hydrothermal fluids. Geochimica et Cosmochimica Acta, 406, 134–147. 10.1016/j.gca.2024.12.019

Lemke, K. H., Rosenbauer, R. J., & Bird, D. K. (2009). Peptide Synthesis in Early Earth Hydrothermal Systems. Astrobiology, 9(2), 141–146. 10.1089/ast.2008.0166

Liu, W., Etschmann, B., Brugger, J., Spiccia, L., Foran, G., & McInnes, B. (2006). UV–Vis spectrophotometric and XAFS studies of ferric chloride complexes in hyper-saline LiCl solutions at 25–90 °C. Chemical Geology, 231(4), 326–349. 10.1016/j.chemgeo.2006.02.005

Liu, W., Borg, S. J., Testemale, D., Etschmann, B., Hazemann, J.-L., & Brugger, J. (2011). Speciation and thermodynamic properties for cobalt chloride complexes in hydrothermal fluids at 35–440 °C and 600 bar: An in-situ XAS study. Geochimica et Cosmochimica Acta, 75(5), 1227–1248. 10.1016/j.gca.2010.12.002

Liu, X., & Xiao, C. (2020). Wolframite solubility and precipitation in hydrothermal fluids: Insight from thermodynamic modeling. Ore Geology Reviews, 117, 103289. 10.1016/j.oregeorev.2019.103289

Liu, X., Xiao, C., & Wang, Y. (2021). The relative solubilities of wolframite and scheelite in hydrothermal fluids: Insights from thermodynamic modeling. Chemical Geology, 584, 120488. 10.1016/j.chemgeo.2021.120488

Liu, X., Yu, P., & Xiao, C. (2023). Tin transport and cassiterite precipitation from hydrothermal fluids. Geoscience Frontiers, 14(6), 101624. 10.1016/j.gsf.2023.101624

Loges, A., Migdisov, A. A., Wagner, T., Williams-Jones, A. E., & Markl, G. (2013). An experimental study of the aqueous solubility and speciation of Y(III) fluoride at temperatures up to 250 °C. Geochimica et Cosmochimica Acta, 123, 403–415. 10.1016/j.gca.2013.07.031

Lowe, A. R., Cox, J. S., & Tremaine, P. R. (2017). Thermodynamics of aqueous adenine: Standard partial molar volumes and heat capacities of adenine, adeninium chloride, and sodium adeninate from *T* = 283.15 K to 363.15 K. The Journal of Chemical Thermodynamics, 112, 129–145. 10.1016/j.jct.2017.04.005

Majzlan, J., Stevens, R., Boerio-Goates, J., Woodfield, B. F., Navrotsky, A., Burns, P. C., et al. (2004). Thermodynamic properties, low-temperature heat-capacity anomalies, and single-crystal X-ray refinement of hydronium jarosite, (H3O)Fe3(SO4)2(OH)6. Physics and Chemistry of Minerals, 31(8), 518–531. 10.1007/s00269-004-0405-z

Majzlan, Juraj, Lang, B. E., Stevens, R., Navrotsky, A., Woodfield, B. F., & Boerio-Goates, J. (2003). Thermodynamics of Fe oxides: Part I. Entropy at standard temperature and pressure and heat capacity of goethite (α-FeOOH), lepidocrocite (γ-FeOOH), and maghemite (γ-Fe2O3). American Mineralogist, 88(5–6), 846–854. 10.2138/am-2003-5-613

Majzlan, Juraj, Grevel, K.-D., & Navrotsky, A. (2003). Thermodynamics of Fe oxides: Part II. Enthalpies of formation and relative stability of goethite (α-FeOOH), lepidocrocite (γ-FeOOH), and maghemite (γ-Fe2O3). American Mineralogist, 88(5–6), 855–859. 10.2138/am-2003-5-614

Majzlan, Juraj, Navrotsky, A., McCleskey, R. B., & Alpers, C. N. (2006). Thermodynamic properties and crystal structure refinement of ferricopiapite, coquimbite, rhomboclase, and Fe2(SO4)3(H2O)5. European Journal of Mineralogy, 175–186. 10.1127/0935-1221/2006/0018-0175

Málek, J., Mitsuhashi, T., Ohashi, N., Taniguchi, Y., Kawaji, H., & Atake, T. (2011). Heat capacity and thermodynamic properties of germanium disulfide at temperatures from *T* = (2 to 1240) K. The Journal of Chemical Thermodynamics, 43(3), 405–409. 10.1016/j.jct.2010.10.011

Marini, L., & Accornero, M. (2007). Prediction of the thermodynamic properties of metal–arsenate and metal–arsenite aqueous complexes to high temperatures and pressures and some geological consequences. Environmental Geology, 52(7), 1343–1363. 10.1007/s00254-006-0578-5

Marini, L., & Accornero, M. (2010). Erratum to: Prediction of the thermodynamic properties of metal–arsenate and metal–arsenite aqueous complexes to high temperatures and pressures and some geological consequences. Environmental Earth Sciences, 59(7), 1601–1606. 10.1007/s12665-009-0369-x

Martín, J., & Delgado Soleri Gil, A. (2010). Ilvaite stability in skarns from the northern contact of the Maladeta batholith, Central Pyrenees (Spain). European Journal of Mineralogy, 363–380. 10.1127/0935-1221/2010/0022-2021

McCollom, T. M., & Shock, E. L. (1997). Geochemical constraints on chemolithoautotrophic metabolism by microorganisms in seafloor hydrothermal systems. Geochimica et Cosmochimica Acta, 61(20), 4375–4391. 10.1016/S0016-7037(97)00241-X

Mercury, L., Vieillard, P., & Tardy, Y. (2001). Thermodynamics of ice polymorphs and ‘ice-like’ water in hydrates and hydroxides. Applied Geochemistry, 16(2), 161–181. 10.1016/S0883-2927(00)00025-1

Migdisov, A., Williams-Jones, A. E., Brugger, J., & Caporuscio, F. A. (2016). Hydrothermal transport, deposition, and fractionation of the REE: Experimental data and thermodynamic calculations. Chemical Geology, 439, 13–42. 10.1016/j.chemgeo.2016.06.005

Migdisov, A., Bastrakov, E., Alcorn, C., Reece, M., Boukhalfa, H., Capporuscio, F. A., & Jove-Colon, C. (2025). A spectroscopic study of the stability of uranyl-carbonate complexes at 25–150 °C and re-visiting the data available for uranyl-chloride, uranyl-sulfate, and uranyl-hydroxide species. Geochimica et Cosmochimica Acta, 406, 326–339. 10.1016/j.gca.2024.04.023

Migdisov, Art. A., Williams-Jones, A. E., & Wagner, T. (2009). An experimental study of the solubility and speciation of the Rare Earth Elements (III) in fluoride- and chloride-bearing aqueous solutions at temperatures up to 300 °C. Geochimica et Cosmochimica Acta, 73(23), 7087–7109. 10.1016/j.gca.2009.08.023

Miron, G. D., Wagner, T., Kulik, D. A., & Heinrich, C. A. (2016). Internally consistent thermodynamic data for aqueous species in the system Na–K–Al–Si–O–H–Cl. Geochimica et Cosmochimica Acta, 187, 41–78. 10.1016/j.gca.2016.04.026

Miron, G. D., Wagner, T., Kulik, D. A., & Lothenbach, B. (2017). An internally consistent thermodynamic dataset for aqueous species in the system Ca-Mg-Na-K-Al-Si-O-H-C-Cl to 800 °C and 5 kbar. American Journal of Science, 317(7). 10.2475/07.2017.01

Morss, L. R., & Konings, R. J. M. (2004). Thermochemistry of Binary Rare Earth Oxides. In G. Adachi, N. Imanaka, & Z. C. Kang (Eds.), Binary Rare Earth Oxides (pp. 163–188). Dordrecht: Springer Netherlands. 10.1007/1-4020-2569-6_7

Murphy, W. M., & Shock, E. L. (1999). Environmental aqueous geochemistry of actinides. Reviews in Mineralogy and Geochemistry, 38(1), 221–253.

Naumov, G. B., Ryzhenko, B. N., Khodakovskiĭ, I. L., Barnes, I., & Speltz, V. (1974). Handbook of thermodynamic data. Menlo Park, Calif., Springfield, Va.: U.S. Geological Survey, Water Resources Division ; [Distributed by] N.T.I.S.

Navrotsky, A., Lee, W., Mielewczyk-Gryn, A., Ushakov, S. V., Anderko, A., Wu, H., & Riman, R. E. (2015). Thermodynamics of solid phases containing rare earth oxides. The Journal of Chemical Thermodynamics, 88, 126–141. 10.1016/j.jct.2015.04.008

Nordstrom, D. K., & Archer, D. G. (2003). Arsenic thermodynamic data and environmental geochemistry. In A. H. Welch & K. G. Stollenwerk (Eds.), Arsenic in Ground Water: Geochemistry and Occurrence (pp. 1–25). Boston, MA: Springer US. 10.1007/0-306-47956-7_1

Oelkers, E. H., & Helgeson, H. C. (1990). Triple-ion anions and polynuclear complexing in supercritical electrolyte solutions. Geochimica et Cosmochimica Acta, 54(3), 727–738. 10.1016/0016-7037(90)90368-U

O’Hare, P. A. G., & Curtiss, L. A. (1995). Thermochemistry of (germanium + sulfur): IV. Critical evaluation of the thermodynamic properties of solid and gaseous germanium(II) sulfide GeS and germanium(IV) disulfide GeS2, and digermanium disulfide Ge2S2(g). Enthalpies of dissociation of bonds in GeS(g), GeS2(g), and Ge2S2(g). The Journal of Chemical Thermodynamics, 27(6), 643–662. 10.1006/jcht.1995.0066

Pankratz, L. B. (1970). Thermodynamic data for silver chloride and silver bromide (Vol. 7430). Pittsburgh, PA: US Bureau of Mines.

Pankratz, L. B., & King, E. G. (1970). High-temperature Enthalpies and Entropies of Chalcopyrite and Bornite. U.S. Department of Interior, Bureau of Mines.

Pankratz, L. B., & Mrazek, R. V. (1982). Thermodynamic properties of elements and oxides. Washington, D.C.: U.S. Dept. of the Interior, Bureau of Mines : For sale by U.S. G.P.O.

Pankratz, L. B., Mah, A. D., & Watson, S. W. (1987). Thermodynamic properties of sulfides. Washington, D.C.: U.S. Dept. of the Interior, Bureau of Mines. Retrieved from http://digicoll.manoa.hawaii.edu/techreports/PDF/USBM-689.pdf

Parker, V. B., & Khodakovskii, I. L. (1995). Thermodynamic Properties of the Aqueous Ions (2+ and 3+) of Iron and the Key Compounds of Iron. Journal of Physical and Chemical Reference Data, 24(5), 1699–1745. 10.1063/1.555964

Perfetti, E., Pokrovski, G. S., Ballerat-Busserolles, K., Majer, V., & Gibert, F. (2008). Densities and heat capacities of aqueous arsenious and arsenic acid solutions to 350 °C and 300 bar, and revised thermodynamic properties of As(OH)3°(aq), AsO(OH)3°(aq) and iron sulfarsenide minerals. Geochimica et Cosmochimica Acta, 72(3), 713–731. 10.1016/j.gca.2007.11.017

Plummer, L. N., & Busenberg, E. (1982). The solubilities of calcite, aragonite and vaterite in CO2-H2O solutions between 0 and 90°C, and an evaluation of the aqueous model for the system CaCO3-CO2-H2O. Geochimica et Cosmochimica Acta, 46(6), 1011–1040. 10.1016/0016-7037(82)90056-4

Plyasunov, A. V., & Shock, E. L. (2001). Correlation strategy for determining the parameters of the revised Helgeson-Kirkham-Flowers model for aqueous nonelectrolytes. Geochimica et Cosmochimica Acta, 65(21), 3879–3900. 10.1016/S0016-7037(01)00678-0

Pokrovski, G. S., & Dubessy, J. (2015). Stability and abundance of the trisulfur radical ion S3− in hydrothermal fluids. Earth and Planetary Science Letters, 411, 298–309. 10.1016/j.epsl.2014.11.035

Pokrovski, G. S., & Schott, J. (1998). Thermodynamic properties of aqueous Ge(IV) hydroxide complexes from 25 to 350°C: implications for the behavior of germanium and the Ge/Si ratio in hydrothermal fluids. Geochimica et Cosmochimica Acta, 62(9), 1631–1642. 10.1016/S0016-7037(98)00081-7

Pokrovski, G. S., Akinfiev, N. N., Borisova, A. Y., Zotov, A. V., & Kouzmanov, K. (2014). Gold speciation and transport in geological fluids: insights from experiments and physical-chemical modelling. Geological Society, London, Special Publications, 402(1), 9–70. 10.1144/SP402.4

Pokrovskii, V. A., & Helgeson, H. C. (1995). Thermodynamic properties of aqueous species and the solubilities of minerals at high pressures and temperatures; the system Al2O3-H2O-NaCl. American Journal of Science, 295(10). 10.2475/ajs.295.10.1255

Pokrovskii, V. A., & Helgeson, H. C. (1997). Thermodynamic properties of aqueous species and the solubilities of minerals at high pressures and temperatures: the system Al2O3-H2O-KOH. Chemical Geology, 137(3), 221–242. 10.1016/S0009-2541(96)00167-2

Popa, K., & Konings, R. J. M. (2006). High-temperature heat capacities of EuPO4 and SmPO4 synthetic monazites. Thermochimica Acta, 445(1), 49–52. 10.1016/j.tca.2006.03.023

Prapaipong, P., Shock, E. L., & Koretsky, C. M. (1999). Metal-organic complexes in geochemical processes: temperature dependence of the standard thermodynamic properties of aqueous complexes between metal cations and dicarboxylate ligands. Geochimica et Cosmochimica Acta, 63(17), 2547–2577. 10.1016/S0016-7037(99)00146-5

Reardon, E. J., & Armstrong, D. K. (1987). Celestite (SrSO4(s)) solubility in water, seawater and NaCl solution. Geochimica et Cosmochimica Acta, 51(1), 63–72. 10.1016/0016-7037(87)90007-X

Richard, L. (2001). Calculation of the standard molal thermodynamic properties as a function of temperature and pressure of some geochemically important organic sulfur compounds§. Geochimica et Cosmochimica Acta, 65(21), 3827–3877. 10.1016/S0016-7037(01)00761-X

Richard, L., & Gaona, X. (2011). Thermodynamic properties of organic iodine compounds. Geochimica et Cosmochimica Acta, 75(22), 7304–7350. 10.1016/j.gca.2011.07.030

Richard, L., & Helgeson, H. C. (1998). Calculation of the thermodynamic properties at elevated temperatures and pressures of saturated and aromatic high molecular weight solid and liquid hydrocarbons in kerogen, bitumen, petroleum, and other organic matter of biogeochemical interest. Geochimica et Cosmochimica Acta, 62(23–24), 3591–3636. 10.1016/S0016-7037(97)00345-1

Robie, R. A., & Hemingway, B. S. (1972). The Heat Capacities at Low-Temperatures and Entropies at 298.15 K of Nesquehonite, MgCO3-3H2O, and Hydromagnesite1. American Mineralogist, 57(11–12), 1768–1781.

Robie, R. A., & Hemingway, B. S. (1995). Thermodynamic properties of minerals and related substances at 298.15 K and 1 bar (10^5^ pascals) pressure and at higher temperatures (No. 2131). Bulletin. U.S. G.P.O. ; For sale by U.S. Geological Survey, Information Services,. 10.3133/b2131

Robie, R. A., Hemingway, B. S., & Fisher, J. R. (1979). Thermodynamic properties of minerals and related substances at 298.15 K and 1 Bar (10^5^ Pascals) pressure and at higher temperatures. U.S. Geological Survey Bulletin, 1452. Retrieved from https://pubs.usgs.gov/bul/1452/report.pdf

Robinson, K. J., Seewald, J. S., Sylva, S. P., Fecteau, K. M., & Shock, E. L. (2024). Thermodynamic property estimations for aqueous primary, secondary, and tertiary alkylamines, benzylamines, and their corresponding aminiums across temperature and pressure are validated by measurements from experiments. Geochimica et Cosmochimica Acta, 372, 62–80. 10.1016/j.gca.2024.03.013

Ruaya, J. R., & Seward, T. M. (1987). The ion-pair constant and other thermodynamic properties of HCl up to 350°C. Geochimica et Cosmochimica Acta, 51(1), 121–130. 10.1016/0016-7037(87)90013-5

Sassani, D. C., & Shock, E. L. (1998). Solubility and transport of platinum-group elements in supercritical fluids: summary and estimates of thermodynamic properties for ruthenium, rhodium, palladium, and platinum solids, aqueous ions, and complexes to 1000°C and 5 kbar. Geochimica et Cosmochimica Acta, 62(15), 2643–2671. 10.1016/S0016-7037(98)00049-0

Schulte, M. (2010). Organic Sulfides in Hydrothermal Solution: Standard Partial Molal Properties and Role in Organic Geochemistry of Hydrothermal Environments. Aquatic Geochemistry, 16(4), 621–637. 10.1007/s10498-010-9102-3

Schulte, M. D., & Rogers, K. L. (2004). Thiols in hydrothermal solution: standard partial molal properties and their role in the organic geochemistry of hydrothermal environments3. Geochimica et Cosmochimica Acta, 68(5), 1087–1097. 10.1016/j.gca.2003.06.001

Schulte, M. D., & Shock, E. L. (1993). Aldehydes in hydrothermal solution: Standard partial molal thermodynamic properties and relative stabilities at high temperatures and pressures. Geochimica et Cosmochimica Acta, 57(16), 3835–3846. 10.1016/0016-7037(93)90337-V

Schulte, M. D., Shock, E. L., & Wood, R. H. (2001). The temperature dependence of the standard-state thermodynamic properties of aqueous nonelectrolytes. Geochimica et Cosmochimica Acta, 65(21), 3919–3930. 10.1016/S0016-7037(01)00717-7

Senoh, H., Ueda, M., Furukawa, N., Inoue, H., & Iwakura, C. (1998). Theoretical evaluation for thermodynamic stability of constituents of Mm-based hydrogen storage alloy in 6 M KOH solution at relatively high temperatures. Journal of Alloys and Compounds, 280(1), 114–124. 10.1016/S0925-8388(98)00739-7

Sharygin, A. V., Grafton, B. K., Xiao, C., Wood, R. H., & Balashov, V. N. (2006). Dissociation constants and speciation in aqueous Li2SO4 and K2SO4 from measurements of electrical conductance to 673 K and 29 MPa. Geochimica et Cosmochimica Acta, 70(20), 5169–5182. 10.1016/j.gca.2006.07.034

Shock, Everetr L., & Koretsky, C. M. (1995). Metal-organic complexes in geochemical processes: Estimation of standard partial molal thermodynamic properties of aqueous complexes between metal cations and monovalent organic acid ligands at high pressures and temperatures. Geochimica et Cosmochimica Acta, 59(8), 1497–1532. 10.1016/0016-7037(95)00058-8

Shock, Everett L. (1992). Stability of peptides in high-temperature aqueous solutions. Geochimica et Cosmochimica Acta, 56(9), 3481–3491. 10.1016/0016-7037(92)90392-V

Shock, Everett L. (1993). Hydrothermal dehydration of aqueous organic compounds. Geochimica et Cosmochimica Acta, 57(14), 3341–3349. 10.1016/0016-7037(93)90542-5

Shock, Everett L. (1995). Organic acids in hydrothermal solutions; standard molal thermodynamic properties of carboxylic acids and estimates of dissociation constants at high temperatures and pressures. American Journal of Science, 295(5), 496–580.

Shock, Everett L., & Helgeson, H. C. (1988). Calculation of the thermodynamic and transport properties of aqueous species at high pressures and temperatures: Correlation algorithms for ionic species and equation of state predictions to 5 kb and 1000°C. Geochimica et Cosmochimica Acta, 52(8), 2009–2036. 10.1016/0016-7037(88)90181-0

Shock, Everett L., & Helgeson, H. C. (1990). Calculation of the thermodynamic and transport properties of aqueous species at high pressures and temperatures: Standard partial molal properties of organic species. Geochimica et Cosmochimica Acta, 54(4), 915–945. 10.1016/0016-7037(90)90429-O

Shock, Everett L., & Koretsky, C. M. (1993). Metal-organic complexes in geochemical processes: Calculation of standard partial molal thermodynamic properties of aqueous acetate complexes at high pressures and temperatures. Geochimica et Cosmochimica Acta, 57(20), 4899–4922. 10.1016/0016-7037(93)90128-J

Shock, Everett L., & McKinnon, W. B. (1993). Hydrothermal Processing of Cometary Volatiles—Applications to Triton. Icarus, 106(2), 464–477. 10.1006/icar.1993.1185

Shock, Everett L, Helgeson, H. C., & Sverjensky, D. A. (1989). Calculation of the thermodynamic and transport properties of aqueous species at high pressures and temperatures: Standard partial molal properties of inorganic neutral species. Geochimica et Cosmochimica Acta, 53(9), 2157–2183. 10.1016/0016-7037(89)90341-4

Shock, Everett L., Sassani, D. C., Willis, M., & Sverjensky, D. A. (1997). Inorganic species in geologic fluids: Correlations among standard molal thermodynamic properties of aqueous ions and hydroxide complexes. Geochimica et Cosmochimica Acta, 61(5), 907–950. 10.1016/S0016-7037(96)00339-0

Shock, Everett L., Sassani, D. C., & Betz, H. (1997). Uranium in geologic fluids: Estimates of standard partial molal properties, oxidation potentials, and hydrolysis constants at high temperatures and pressures. Geochimica et Cosmochimica Acta, 61(20), 4245–4266. 10.1016/S0016-7037(97)00240-8

Shvarov, Y. V., & Bastrakov, E. N. (1999). HCh: a software package for geochemical equilibrium modelling. User’s Guide. Record 1999/025. Australian Geological Survey Organisation. Retrieved from **Error! Hyperlink reference not valid**.

Shvarov, Yu. V. (2008). HCh: New potentialities for the thermodynamic simulation of geochemical systems offered by windows. Geochemistry International, 46(8), 834–839. 10.1134/S0016702908080089

Shvedov, D., & Tremaine, P. R. (1997). Thermodynamic Properties of Aqueous Dimethylamine and Dimethylammonium Chloride at Temperatures from 283 K to 523 K: Apparent Molar Volumes, Heat Capacities, and Temperature Dependence of Ionization. Journal of Solution Chemistry, 26(12), 1113–1143. 10.1023/A:1022977006327

St Clair, B., Pottenger, J., Debes, R., Hanselmann, K., & Shock, E. (2019). Distinguishing Biotic and Abiotic Iron Oxidation at Low Temperatures. ACS Earth and Space Chemistry, 3(6), 905–921. 10.1021/acsearthspacechem.9b00016

Stefánsson, A. (2001). Dissolution of primary minerals of basalt in natural waters: I. Calculation of mineral solubilities from 0°C to 350°C. Chemical Geology, 172(3), 225–250. 10.1016/S0009-2541(00)00263-1

Stefánsson, A., Bénézeth, P., & Schott, J. (2013). Carbonic acid ionization and the stability of sodium bicarbonate and carbonate ion pairs to 200 °C – A potentiometric and spectrophotometric study. Geochimica et Cosmochimica Acta, 120, 600–611. 10.1016/j.gca.2013.04.023

Stefánsson, A., Bénézeth, P., & Schott, J. (2014). Potentiometric and spectrophotometric study of the stability of magnesium carbonate and bicarbonate ion pairs to 150 °C and aqueous inorganic carbon speciation and magnesite solubility. Geochimica et Cosmochimica Acta, 138, 21–31. 10.1016/j.gca.2014.04.008

Stoffregen, R. E., Alpers, C. N., & Jambor, J. L. (2000). Alunite-Jarosite Crystallography, Thermodynamics, and Geochronology. Reviews in Mineralogy and Geochemistry, 40(1), 453–479. 10.2138/rmg.2000.40.9

Sverjensky, D. A., Shock, E. L., & Helgeson, H. C. (1997). Prediction of the thermodynamic properties of aqueous metal complexes to 1000°C and 5 kb. Geochimica et Cosmochimica Acta, 61(7), 1359–1412. 10.1016/S0016-7037(97)00009-4

Sverjensky, Dimitri A., Hemley, J. J., & D’angelo, W. M. (1991). Thermodynamic assessment of hydrothermal alkali feldspar-mica-aluminosilicate equilibria. Geochimica et Cosmochimica Acta, 55(4), 989–1004. 10.1016/0016-7037(91)90157-Z

Sverjensky, Dimitri A., Harrison, B., & Azzolini, D. (2014). Water in the deep Earth: The dielectric constant and the solubilities of quartz and corundum to 60 kb and 1200 °C. Geochimica et Cosmochimica Acta, 129, 125–145. 10.1016/j.gca.2013.12.019

Tagirov, B., & Schott, J. (2001). Aluminum speciation in crustal fluids revisited. Geochimica et Cosmochimica Acta, 65(21), 3965–3992. 10.1016/S0016-7037(01)00705-0

Tagirov, B. R., Zotov, A. V., & Akinfiev, N. N. (1997). Experimental study of dissociation of HCl from 350 to 500°C and from 500 to 2500 bars: Thermodynamic properties of HCl°(aq). Geochimica et Cosmochimica Acta, 61(20), 4267–4280. 10.1016/S0016-7037(97)00274-3

Tagirov, B. R., Baranova, N. N., & Bychkova, Ya. V. (2015). Thermodynamic properties of platinum chloride complexes in aqueous solutions: Derivation of consistent parameters from literature data and experiments on Pt(cr) solubility at 400–475°C and 1 kbar. Geochemistry International, 53(4), 327–340. 10.1134/S0016702915040084

Tagirov, B. R., Akinfiev, N. N., & Zotov, A. V. (2024). Gold(I) Complexation in Chloride Hydrothermal Fluids. Geology of Ore Deposits, 66(5), 581–597. 10.1134/S1075701524600403

Tagirov, Boris R., Baranova, N. N., Zotov, A. V., Akinfiev, N. N., Polotnyanko, N. A., Shikinal, N. D., et al. (2013). The speciation and transport of palladium in hydrothermal fluids: Experimental modeling and thermodynamic constraints. Geochimica et Cosmochimica Acta, 117, 348–373. 10.1016/j.gca.2013.03.047

Tagirov, Boris R., Akinfiev, N. N., Tarnopolskaia, M. E., Nikolaeva, I. Yu., Zlivko, I. Yu., Volchenkova, V. A., et al. (2025). Gold in sulfide fluids revisited. Geochimica et Cosmochimica Acta, 406, 285–304. 10.1016/j.gca.2024.08.022

Tardy, Y., Schaul, R., & Duplay, J. (1997). Domaines de stabilité thermodynamiques des humus, de la microflore et des plantes. Comptes Rendus de l’Académie Des Sciences - Series IIA - Earth and Planetary Science, 324(12), 969–976. 10.1016/S1251-8050(97)83981-X

Trofimov, N. D., Tagirov, B. R., Akinfiev, N. N., Reukov, V. L., Nickolsky, M. S., Nikolaeva, I. Yu., et al. (2023). Chalcocite Cu2S solubility in aqueous sulfide and chloride fluids. Thermodynamic properties of copper(I) aqueous species and copper transport in hydrothermal systems. Chemical Geology, 625, 121413. 10.1016/j.chemgeo.2023.121413

Vidal, O., Goffé, B., & Theye, T. (1992). Experimental study of the stability of sudoite and magnesiocarpholite and calculation of a new petrogenetic grid for the system FeO–MgO–Al2O3–SiO2–H2O. Journal of Metamorphic Geology, 10(5), 603–614. 10.1111/j.1525-1314.1992.tb00109.x

Vidal, Olivier, Parra, T., & Trotet, F. (2001). A Thermodynamic Model for Fe-Mg Aluminous Chlorite Using Data from Phase Equilibrium Experiments and Natural Pelitic Assemblages in the 100° to 600°c, 1 to 25 kb Range. American Journal of Science, 301(6). 10.2475/ajs.301.6.557

Vidal, Olivier, Parra, T., & Vieillard, P. (2005). Thermodynamic properties of the Tschermak solid solution in Fe-chlorite: Application to natural examples and possible role of oxidation. American Mineralogist, 90(2–3), 347–358. 10.2138/am.2005.1554

Von Der Heyden, B. P., Dick, J., Rosenfels, R. C., Carlton, L., Lilova, K., Navrotsky, A., et al. (2024). Growth and Stability of Stratiform Carrollite (CuCo2S4) in the Tenke-Fungurume Ore District, Central African Copperbelt. The Canadian Journal of Mineralogy and Petrology, 62(1), 77–93. 10.3749/2300028

Wagman, D. D., Evans, W. H., Parker, V. B., Schumm, R. H., Halow, I., Bailey, S. M., et al. (1982). The NBS tables of chemical thermodynamic properties: Selected values for inorganic and C1 and C2 organic substances in SI units. Journal of Physical and Chemical Reference Data, 11(2). Retrieved from https://srd.nist.gov/JPCRD/jpcrdS2Vol11.pdf

Williams-Jones, A. E., & Vasyukova, O. V. (2022). Constraints on the Genesis of Cobalt Deposits: Part I. Theoretical Considerations. Economic Geology, 117(3), 513–528. 10.5382/econgeo.4895

Wood, S. A., & Samson, I. M. (2000). The Hydrothermal Geochemistry of Tungsten in Granitoid Environments: I. Relative Solubilities of Ferberite and Scheelite as a Function of T, P, pH, and mNaCl. Economic Geology, 95(1), 143–182. 10.2113/gsecongeo.95.1.143

Zhu, C., & Sverjensky, D. A. (1992). F-Cl-OH partitioning between biotite and apatite. Geochimica et Cosmochimica Acta, 56(9), 3435–3467. 10.1016/0016-7037(92)90390-5

Zhu, Y., Zhang, X., Xie, Q., Chen, Y., Wang, D., Liang, Y., & Lu, J. (2005). Solubility and stability of barium arsenate and barium hydrogen arsenate at 25 °C. Journal of Hazardous Materials, 120(1), 37–44. 10.1016/j.jhazmat.2004.12.025

Ziemer, S. P., & Woolley, E. M. (2007). Thermodynamics of the first and second proton dissociations from aqueous l-aspartic acid and l-glutamic acid at temperatures from (278.15 to 393.15) K and at the pressure 0.35 MPa: Apparent molar heat capacities and apparent molar volumes of zwitterionic, protonated cationic, and deprotonated anionic forms at molalities from (0.002 to 1.0) mol · kg−1. The Journal of Chemical Thermodynamics, 39(4), 645–666. 10.1016/j.jct.2006.08.008

Zimmer, K., Zhang, Y., Lu, P., Chen, Y., Zhang, G., Dalkilic, M., & Zhu, C. (2016). SUPCRTBL: A revised and extended thermodynamic dataset and software package of SUPCRT92. Computers & Geosciences, 90, 97–111. 10.1016/j.cageo.2016.02.013

